# Prolonged Crowding Initiates Tumor Invasion with Mechanomemory by Pressure-Sensation of Membrane Domains

**DOI:** 10.1101/2024.07.28.605413

**Authors:** Xinbin Zhao, Min Tan, Long Li, Xubo Lin, Zekun Li, Xiaohuan Wang, Bingqi Song, Zheng Guo, Tailin Chen, Sen Hou, Jiehou Fan, Shijiang Wang, Yun Zhang, Yubo Fan, Jing Du

## Abstract

During the progression from epithelial neoplasms to invasive carcinoma, cells are subjected to prolonged confinement. However, the response of cancer cells to such mild yet sustained compressive pressure during the initial stages of tumor invasion remain poorly understood. Here, using a spontaneous crowding model to recapitulate the progressive compressive stress caused by cell proliferation, we demonstrated that prolonged crowding alone is sufficient to induce the acquisition of an invasive phenotype and associated gene expression patterns in cancer cells. This invasiveness persisted even after cells were removed from the crowded environment, a phenomenon mediated by mechanomemory. By combining genetic manipulations, mechanical modeling, and biophysical measurements, we revealed that the disaggregation of membrane domains**—**driven by a nanoscale smooth-corrugated topography transition of plasma membranes induced by Laplace pressure under crowded conditions**—**is essential for initiating cancer cell invasion. Inhibiting membrane domains disaggregation through membrane-to-cortex-attachment effectively suppresses cancer cell invasion in both cellular crowding models and mouse xenograft models. This study underscores the critical role of tissue-scale mechanics in regulating the biophysics of mechanosensitive membrane domains during the early stages of tumor invasion.

## INTRODUCTION

Mechanical force modulates embryonic development, influences tissue homeostasis, and contributes to the development of many diseases including cancer^1–3^. The primary tumor microenvironment (TME) is characterized by a diverse array of harsh mechanical cues, including increased matrix stiffness and solid stress^4^. Due to cell growth and surrounding pressures, solid stress is the force transmitted through the elastic solid phase of the tissue and can generate tensile stress and compressive stress^5^. During the development from epithelial neoplasms to invasive carcinoma, uncontrolled growth and proliferation of cancer cells pushes and displaces the surrounding normal tissue, which in turn constrains tumor expansion and results in crowded tissues and generates compressive mechanical stress within solid tumor and surrounding extracellular matrix (ECM)^6–8^.

Recent studies in mouse models of skin tumor reveal that constraining forces from overlying suprabasal cancer cells and underlying ECM shape tissue architecture and affect tumor invasion^9,10^. Strong confinement has been shown to drive a fast amoeboid migration in mesenchymal cells and embryonic progenitor cells^11,12^. The cell nucleus is able to sense constraining forces and responds to them by switching to a rapid migratory phenotypic state that enables cancer cells to squeeze out from compressive conditions^13,14^. Strong confinement during cell invasion also causes nuclear deformation, which results in localized nuclear envelope rupture and DNA damage, and promotes invasive phenotype of MCF10DCIS.com cells^15^. At present, experimental methods used to study the invasion of cancer cells in response to physical confinement mainly subject cells to transient and strong force by *in vitro* compression device or microfabricated duct-on-a-chip which mimicked the invasive process *per se* of cancer cells through a narrow channel^15,16^. However, the mechanical adaptability of cancer cells to the mild and prolonged compression under crowded conditions and their function in the initiation of tumor invasion remain unclear.

Emerging evidence has revealed the ability of cells to remember the mechanical stimuli after the cessation of force, which is termed “mechanomemory”^17^. Mechanomemory has been studied in mesenchymal stem cells and epithelial cells, mostly with high matrix stiffness as a physical stimulus. However, the investigation of cell mechanomemory in the cancer field is currently in its infancy^18,19^. Whether prolonged exposure to compressive stress in the crowded TME could imprint this mechanomemory in cancer cells has not been studied.

In this study, we found that prolonged crowding initiates invasion with mechanomemory in cancer cells by using a spontaneous crowding model composed of a freely growing monoclonal cell sheet. Combining genetic manipulations, biophysical measurements, and mechanical modeling, we revealed that the disaggregation of membrane domains sensitive to prolonged crowding drives cancer cell invasiveness. Membrane domains disaggregation is induced by a nanoscale smooth-corrugated topography transition (nSCTT) of plasma membranes under Laplace pressure. Finally, we demonstrated that enhancing the aggregation of membrane domains by suppressing the nSCTT through membrane-to-cortex-attachment (MCA) inhibits cancer cell invasiveness in cellular crowding models and mouse xenograft models.

## RESULTS

### Prolonged crowding initiates cancer cell invasion

To investigate the role of prolonged crowding in tumor invasion, we employed a spontaneous crowding model^20^ using HeLa cells, which were allowed to freely grow as a monoclonal cell sheet for 14 days (Fig. 1a). According to our previous study, this cell sheet spontaneously developed a progressive crowding gradient radiating from the central region to the periphery driven by interfacial shear stress between the cell sheet and the ECM^21^. The extent of crowding was quantified using a metric termed “crowding strain”, calculated as (*A_0_* – *A_n_*) /*A_0_*, where *A_0_* represents the nuclear area of cells in sparse culture, and *A_n_* denotes the nuclear area of cells within the cell sheet (Fig. 1b). Based on crowding strain values, the cell sheet was segmented into two regions: an uncrowded region (crowding strain ranging from -0.3 to 0) and a crowded region (crowding strain from 0 to 0.3) (Fig. 1c). Using this model, cells in the crowded region were subjected to long-term compressive stress induced by prolonged crowding for 7-14 days. Transwell invasion assays revealed that crowded cells exhibited significantly enhanced invasiveness compared to uncrowded cells (Fig. 1d and e). These findings were corroborated in other cancer cell lines, including MC38 (a murine colorectal cancer cell line) and A431 (a human skin cancer cell line) (Fig. 1d and e). Furthermore, time-lapse imaging of NLS-GFP^+^ HeLa cell sheets demonstrated that cell invasiveness originated predominantly from crowded regions (Fig. S1a and b), providing direct evidence that prolonged crowding initiates cancer cell invasion.

**Figure 1.**
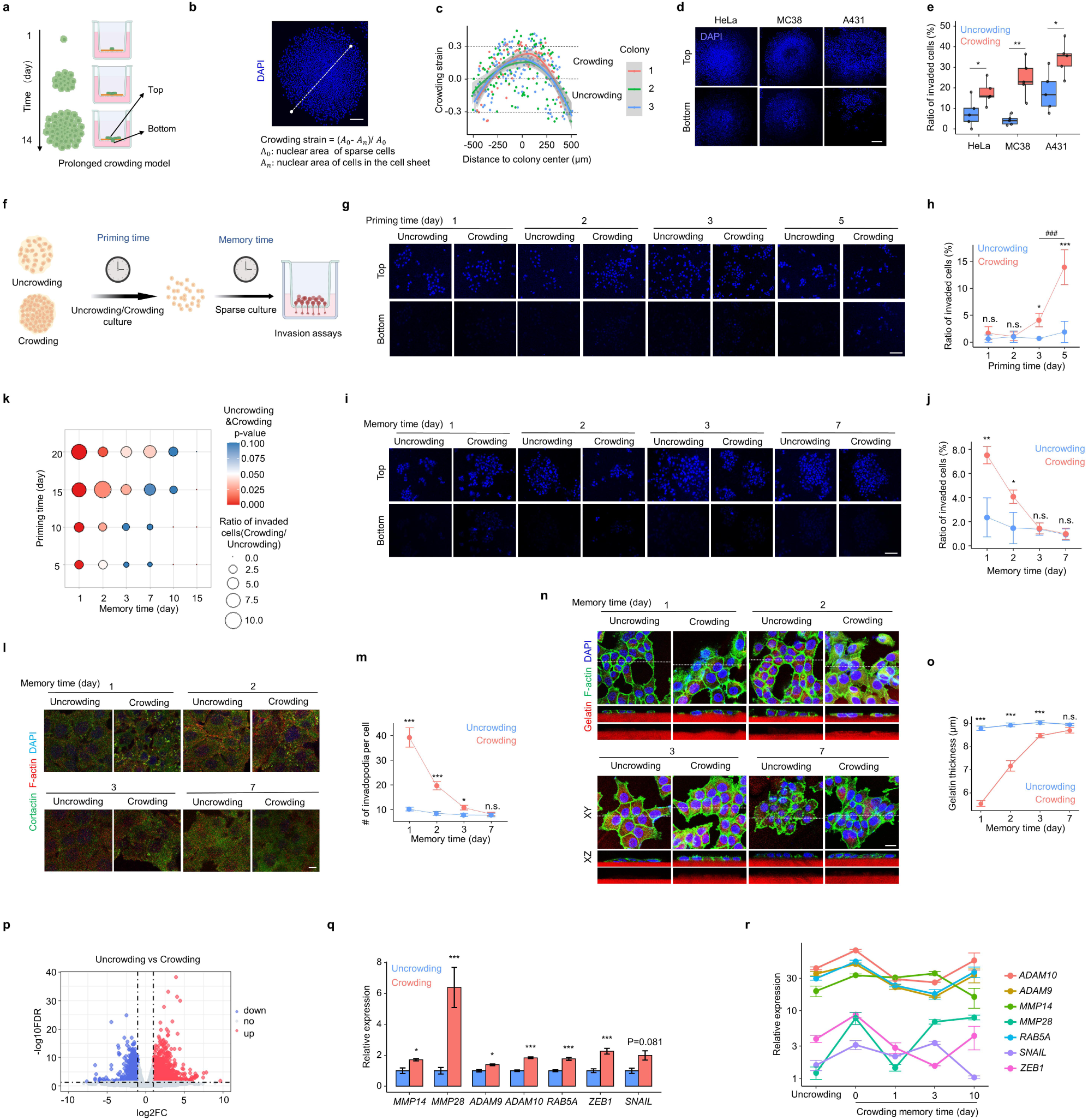
Prolonged crowding initiates cancer cell invasion with mechanomemory. (**a**) Schematic diagram of prolonged crowding-induced cell invasiveness model. (**b**) Quantitation of the individual cell nucleus area as crowding strain along the lines in the panel. Scale bar: 200 μm. (**c**) The statistical analysis of crowding strain per cell along the panel lines (b). The cell sheet was segmented into crowding (crowding strain from 0 to 0.3) and uncrowding (crowding strain from -0.3 to 0). (**d**) Transwell matrigel invasion assay of a growing monoclonal HeLa, MC38, and A431 cell sheets after culture for 14 days. Representative DAPI images of cells that accumulated on the top (uninvaded) and bottom (invaded) surface of the insert membranes. Scale bar: 200 μm. (**e**) The ratio of invaded cells in uncrowded and crowded HeLa, MC38, and A431 cells. (**f**) Schematic illustration of the experimental procedure to test the invasion of HeLa cells after priming time for uncrowding and crowding culture and memory time for sparse culture. (**g**) Transwell matrigel invasion assay of HeLa cells after experiencing uncrowding and crowding for 1, 2, 3, 5 days (priming time), subsequently collected and transferred to transwell chamber for 48 hrs. Representative DAPI images of cells that accumulated on the top and bottom surface of the insert membranes. Scale bar: 100 μm. (**h**) The ratio of invaded cells in uncrowded and crowded HeLa cells in panel (g). (**i**) HeLa cells were grown dishes in crowding or uncrowding culture for 5 days, transferred to new dishes for 1, 2, 3, 7 days (memory time), subsequently collected and transferred to transwell chamber for 48 hrs. Representative DAPI images of cells that accumulated on the top and bottom surface of the insert membranes. Scale bar: 100 μm. (**j**) The ratio of invaded cells in uncrowded and crowded HeLa cells in panel (i). (**k**) The correlation graph between priming time and memory time in cell culture for invasion analysis. (**l**) HeLa cells were grown dishes in crowding or uncrowding culture for 5 days, transferred to matrigel-coated dishes for 1, 2, 3, 7 days. Representative XY slice image of cortactin distribution at the basal membrane in uncrowding/crowding groups. Scale bar: 10 μm. (**m**) Quantification of invadopodia number per cell with corresponding memory time. (**n**) HeLa cells were grown dishes in crowding or uncrowding culture for 5 days, transferred to Cy5-conjugated gelatin hydrogels for 1, 2, 3, 7 days. Representative XY slice image of F-actin staining in uncrowding/crowding groups. Scale bar: 20 μm. (**o**) Quantification of correlation between Cy5-conjugated gelatin thickness underneath cells and its memory time. (**p**) The volcano plot shows deferentially expressed genes between uncrowding and crowding. Each dot represents a gene; genes in red are up, genes in blue are down-regulated. (**q**) Boxplots showing mRNA expression data for *MMP14*, *MMP28*, *ADAM9*, *ADAM10*, *RAB5A*, *ZEB1*, and *SNAIL* in uncrowded and crowded HeLa cells as determined by RNA seq data respectively. (**r**) Relative expression of *MMP14*, *MMP28*, *ADAM9*, *ADAM10*, *RAB5A*, *ZEB1*, and *SNAIL* in HeLa cells after experiencing uncrowding or crowding for 0, 1, 3, 10 days (memory time). Schematic images (c, f) were created with BioRender.com. Data are presented as mean ± SEM; *p < 0.05, **p < 0.01, ***p < 0.005, not significant (n.s.); two-tailed unpaired t-test.

Given that the invasion process is known to require degradation of the surrounding ECM^22^, we next investigated whether prolonged crowding influences ECM degradation. To this end, we utilized Cy5-conjugated gelatin hydrogels coated on culture dishes. Vertical cross-sectional (XZ-plane) imaging revealed that the thickness of the ECM gradually decreased with increasing crowding strain (Fig. S1c and d), demonstrating that prolonged crowding enhances the capacity of HeLa cells to degrade the ECM. Furthermore, immunofluorescence staining of cortactin, an actin-bundling protein enriched in invadopodia of tumor cells^23,24^, showed that the number of invadopodia per cell progressively increased with higher crowding strain in HeLa cells cultured on matrigel-coated culture dishes (Fig. S1e and f). Taken together, these results suggest that prolonged crowding is sufficient to drive the initiation of cancer cell invasion.

### Prolonged crowding-initiated cancer cell invasion is retained via mechanomemory

Recent studies have demonstrated the ability of matrix stiffness to imprint mechanomemory onto the cells after cessation of a force^25,26^. We next investigated whether cells exposed to prolonged crowding retain their invasive properties after leaving crowded regions. HeLa cells were first cultured under crowded or uncrowded conditions for varying durations (priming time). These primed cells were then transferred to sparse culture conditions (crowding strain ≈ 0) for different periods (memory time) before being subjected to invasion assays, such as Transwell assay (Fig. 1f). Cells primed in crowded cultures for 3 and 5 days exhibited significantly elevated invasiveness compared to those primed for 1 and 2 days (Fig. 1g and h). Furthermore, cells primed for 5 days displayed greater invasiveness than those primed for 3 days (Fig. 1h), indicating a priming-time-dependent mechanomemory that governs the retention of invasiveness.

We next investigated the duration of crowding-induced mechanomemory by priming cells for 5 days in crowded or uncrowded cultures, followed by transfer to sparse culture for 1, 2, 3, or 7 days (Fig. 1i). Transwell invasion assays revealed that crowded cells retained significantly higher invasiveness compared to uncrowded cells after 1 and 2 days of sparse culture (Fig. 1i and j). To further explore the relationship between priming duration and mechanomemory, we extended the priming time in crowded cultures up to 20 days. Transwell assays demonstrated that the duration of mechanomemory increased with priming time, with cells primed for 20 days retaining enhanced invasiveness even after 7 days in sparse culture (Fig. 1k). This retention of invasiveness via mechanomemory was consistently observed in A431 cells (Fig. S2). Additionally, crowding-primed cells (5 days) exhibited a significantly higher number of invadopodia per cell compared to uncrowded cells after 1, 2, and 3 days in sparse culture (Fig. 1l and m). Notably, mechanomemory was also evident in the ECM degradation ability of crowding-primed cells (Fig. 1n and o). Together, these results demonstrate that prolonged crowding not only initiates cancer cell invasion but also imprints a duration-dependent mechanomemory, sustaining the invasive phenotype even after the removal of crowding stress.

To further characterize gene expression regulated by prolonged crowding, we performed RNA sequencing of crowding-primed cells (20 days) followed by sparse culture for varying durations (0, 1, 3, or 10 days). We identified 1,990 up-regulated and 1,408 down-regulated genes that were significantly altered in crowded cells compared to uncrowded cells without a sparse culture step (Fig. 1p). Among these, genes involved in ECM degradation, such as *MMP28*, *MMP14, ADAM9,* and *ADAM10*^27^, as well as *RAB5A*, as a micro-invasive marker, were upregulated in crowded cells compared to uncrowded cells (Fig. 1q). These findings align with the enhanced invasiveness observed in crowded cells.

Additionally, we detected significantly elevated levels of epithelial-mesenchymal transition (EMT)-inducing transcription factors, including *SNAIL* and *ZEB1*, in crowded cells (Fig. 1q), indicating a transition towards a more mesenchymal-like state. Notably, the increased expression levels of *MMP14, MMP28*, and *SNAIL* in crowded cells were sustained for at least 3 days in sparse cultures (Fig. 1r), consistent with the retention of mechanomemory. These results demonstrate that prolonged crowding enhances the expression of genes associated with invasiveness and retains these expression patterns through mechanomemory. Collectively, these findings establish that prolonged crowding not only initiates cancer cell invasion but also imprints a mechanical memory that sustains the invasive phenotype even after the removal of crowding stress.

### Prolonged crowding initiates cancer cell invasion by disrupting the aggregation of membrane domains

To elucidate the mechanism underlying the invasion initiated by prolonged crowding, we further analyzed RNA-seq data and correlated gene expression changes with memory time using mFuzz. Our analysis identified 187 genes in Cluster 2 whose expression trends correlated with memory time, showing a recovery in crowded cells after 1, 3, and 10 days of sparse culture compared to uncrowded cells (Fig. S3). Gene Ontology (GO) cellular component enrichment analysis of Cluster 2 genes revealed that most were associated with the cell membrane or membrane domains (Fig. 2a). Previous studies have established that plasma membranes contain numerous lipid microdomains enriched in cholesterol and sphingolipids^28,29^. These microdomains, termed lipid rafts, include subtypes such as flotillin-rich planar lipid rafts and caveolin-rich caveolae^28^. Lipid rafts play critical roles in various physiological and pathological processes by aggregating into larger platforms that regulate signaling regulation^28–30^. To experimentally assess whether membrane domains are affected by prolonged crowding, we monitored the abundance of membrane domains in the cellular crowding model using Cy5-conjugated cholera toxin subunit B (CTxB), which specifically binds to ganglioside GM1, a common ganglioside enriched in membrane domains^31^. Fluorescence imaging revealed that the extent of CTxB-labeled membrane domains at the apical plasma membrane was significantly reduced in crowded cells compared to uncrowded cells (Fig. 2b and c). Specifically, both the fluorescence intensity of CTxB and the number of CTxB clusters at the apical plasma membrane per cell exhibited a crowding-strain-dependent reduction when the crowding strain exceeded 0 (Fig. 2d-f). Further, immunofluorescence staining against Flotillin-1, Caveolin-1, and Cavin-1 also showed significant crowding-strain-dependent decreases in signal intensity (Fig. 2g-j and Fig. S4). These results demonstrate that the aggregation of membrane domains at the apical plasma membrane in cancer cells is disrupted by prolonged crowding.

**Figure 2.**
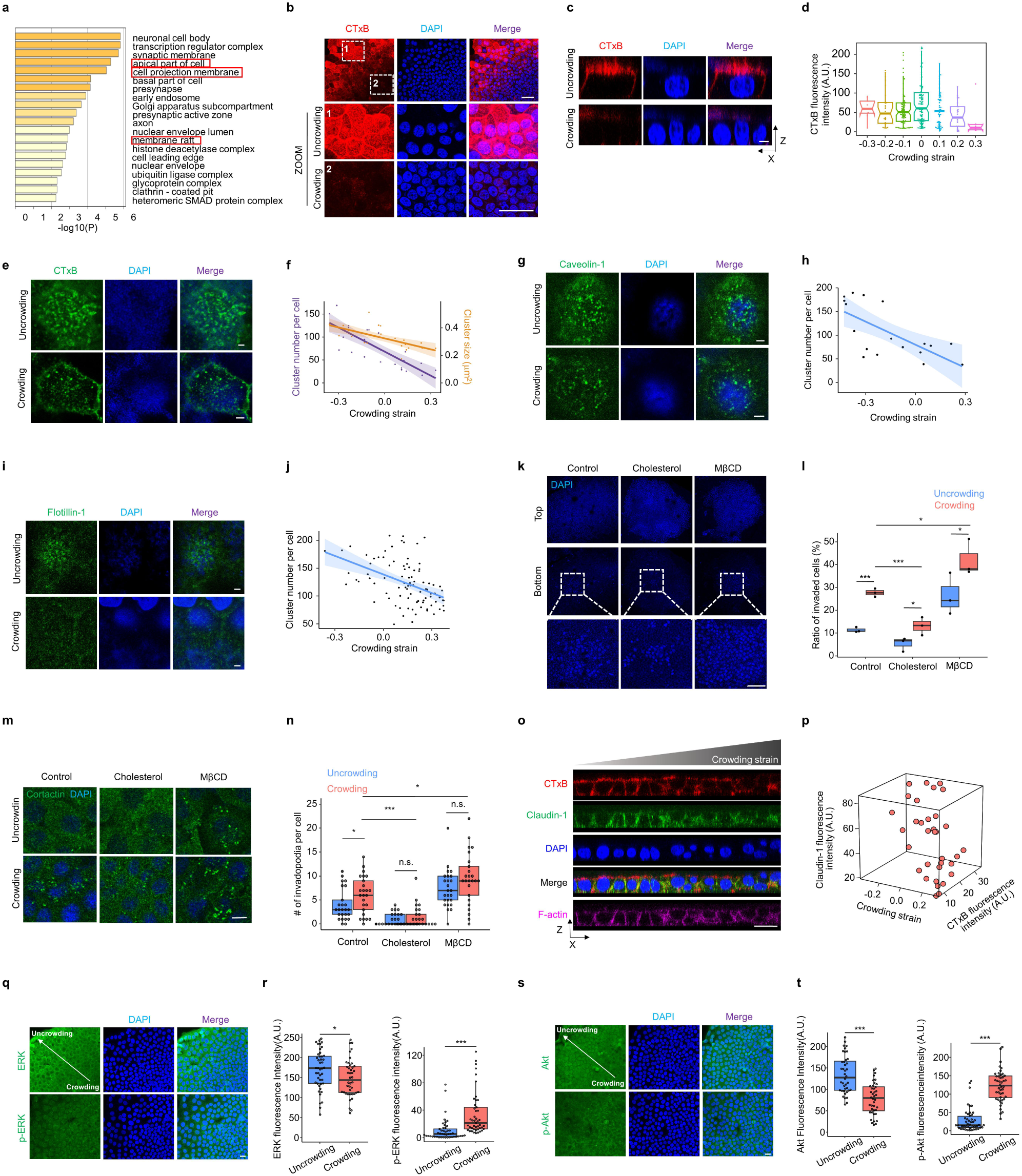
Prolonged crowding promotes cancer cell invasiveness by disrupting the aggregation of membrane domains. (**a**) The Gene Ontology (GO) cellular components enriched by Metascape from RNA sequencing data. (**b**) Representative XY confocal images for CTxB staining in uncrowded and crowded HeLa cells. Scale bar: 50 μm. (**c**) Reconstructed XZ confocal images for CTxB staining at the apical membrane of uncrowded and crowded HeLa cells. Scale bar: 5 μm. (**d**) Quantitation of correlation between CTxB fluorescence intensity per cell and its crowding strain. (**e**) Representative XY confocal images for CTxB staining at the apical membrane of uncrowded and crowded HeLa cells. Scale bar: 2 μm. (**f**) Quantitation of correlation between the number and size of CTxB clusters per cell and its crowding strain. (**g**) Representative XY confocal images showing immunostaining for Caveolin-1 at the apical membrane of uncrowded and crowded HeLa cells. Scale bar: 2 μm. (**h**) Quantitation of correlation between the number of Caveolin-1 clusters per cell and its crowding strain. (**i**) Representative XY confocal images showing immunostaining for Flotillin-1 at the apical membrane of uncrowded and crowded HeLa cells. Scale bar: 2 μm. (**j**) Quantitation of correlation between the number of Flotillin-1 clusters per cell and its crowding strain. (**k**) Representative XY slice images of cholesterol- and MβCD-treated HeLa cell sheets by Transwell invasion assay. Scale bar: 100 μm. (**l**) Quantification of invaded cell number in panel (k). (**m**) Representative XY slice images showing immunostaining for cortactin in uncrowded and crowded cells treated with cholesterol and MβCD. Scale bar: 10 μm. (**n**) Quantification of invadopodia density per cell in panel (m). (**o**) Reconstructed XZ confocal images showing the localization of CTxB and Claudin-1 in the regions with different crowding strain. Size bars: 20 μm. (**p**) Quantitation of correlation between Claudin-1 fluorescence intensity and CTxB fluorescence intensity in differently crowded cells. (**q**) Representative XY confocal images showing immunostaining for ERK and p-ERK in uncrowded and crowded HeLa cells. Scale bar: 2 μm. (**r**) Quantification of fluorescence intensity of ERK and p-ERK per cell in panel (q). (**s**) Representative XY confocal images showing immunostaining for Akt and p-Akt in uncrowded and crowded HeLa cells. Scale bar: 20 μm. (**t**) Quantification of fluorescence intensity of Akt and p-Akt per cell in panel (s). Data are presented as *p < 0.05, ***p < 0.005; two-tailed unpaired t-test.

To investigate whether the disrupted aggregation of membrane domains contributes to initiation of cancer cell invasion under prolonged crowding, we modulated membrane domains by enriching or depleting membrane cholesterol content. This was achieved through the addition of exogenous cholesterol or methyl-β-cyclodextrin (MβCD)^32^, respectively, in the cellular crowding model. Transwell invasion assays revealed that cholesterol enrichment significantly inhibited cell invasion compared to the control group, whereas cholesterol depletion by MβCD significantly enhanced the invasiveness in crowded cells (Fig. 2k, l). Notably, both exogenous cholesterol addition and MβCD treatment reduced the difference in invasiveness between crowded and uncrowded cells (Fig. 2l). Furthermore, exogenous cholesterol addition decreased the number of invadopodia per cell, while MβCD treatment increased it, effectively eliminating the difference between crowded and uncrowded cells (Fig. 2m, n). These findings collectively demonstrate that prolonged crowding initiates cancer cell invasion by disrupting the aggregation of membrane domains.

Extensive studies have established a strong link between carcinoma invasion and the loss of epithelial integrity, particularly through the disruption of cell-cell contacts^33^. Cholesterol-rich membrane domains at apical junctions are known to be essential for tight junction formation^34^. In our study, using an antibody against claudin-1, a key component of tight junction^35^, we observed a progressive decline in tight junction integrity as crowding strain increased in the cellular crowding model (Fig. 2o, p). Furthermore, the fluorescence intensity of CTxB, which labels membrane domains, showed a positively correlated with Claudin-1 intensity (Fig. 2o, p). These results indicate that prolonged crowding initiates cancer cell invasion by disrupting epithelial integrity, a process mediated by the disaggregation of membrane domains.

Numerous studies have shown that PI3K/AKT and MAPK/ERK pathways are associated with tumor progression such as invasion and display activating genetic alterations in more than 40% of primary tumors^36–38^. Next, we wondered whether these pathways participate in tumor invasion initiated by prolonged crowding. Immunofluorescence staining revealed that the phosphorylation of Akt and ERK was significantly upregulated in crowded cells versus uncrowded cells, implying that prolonged crowding activated these signaling pathways (Fig. 2q-t). Together, these findings demonstrate that the activation of Akt and ERK, triggered by prolonged crowding, is implicated in the the initiation of cancer cell invasion.

### Prolonged crowding triggers a nanoscale smooth-corrugated topography transition of plasma membranes

Given that membrane domain availability is regulated by lipid metabolism, we next evaluated the levels of cholesterol and GM1 ganglioside in cells exposed to prolonged crowding^28,31^. However, fluorescent filipin III staining revealed no significant differences in free cholesterol levels between uncrowded cells and crowded cells (Fig. S5a, b). Similarly, immunofluorescence staining using an antibody against GM1 ganglioside indicated that its levels at the apical plasma membrane remained unchanged under different crowding strains (Fig. S5c, d). These findings suggest that prolonged crowding does not significantly alter the levels of main components of membrane domains.

Recent studies reported that the topographic configuration of cellular membranes plays a critical role in regulating membrane domains^39^. To further investigate the mechanism underlying the disaggregation of membrane domains, we first analyzed the topology of plasma membranes. In the cellular crowding model, we observed significant deformation of plasma membranes in crowded cells compared to uncrowded cells (Fig. 3a and b). Furthermore, the intensity of CTxB-labeled membrane domains at the apical plasma membrane progressively decreased as membrane curvature increased (Fig. 3c). We further examined plasma membrane topology in a subcutaneous nude mouse xenograft model using EYFP-mem^+^ HeLa cells. This revealed pronounced plasma membrane deformation in cells within micro-invasive foci exposed to crowding (Fig. 3d-f). Consistent plasma membrane deformation was also observed in human skin cancer tissue (Fig. 3g, h), as well as in cervical and breast cancer tissues (Fig. S6). These results demonstrate that severe plasma membrane deformation, associated with membrane domain disaggregation, occurs in cells exposed to crowded conditions.

**Figure 3.**
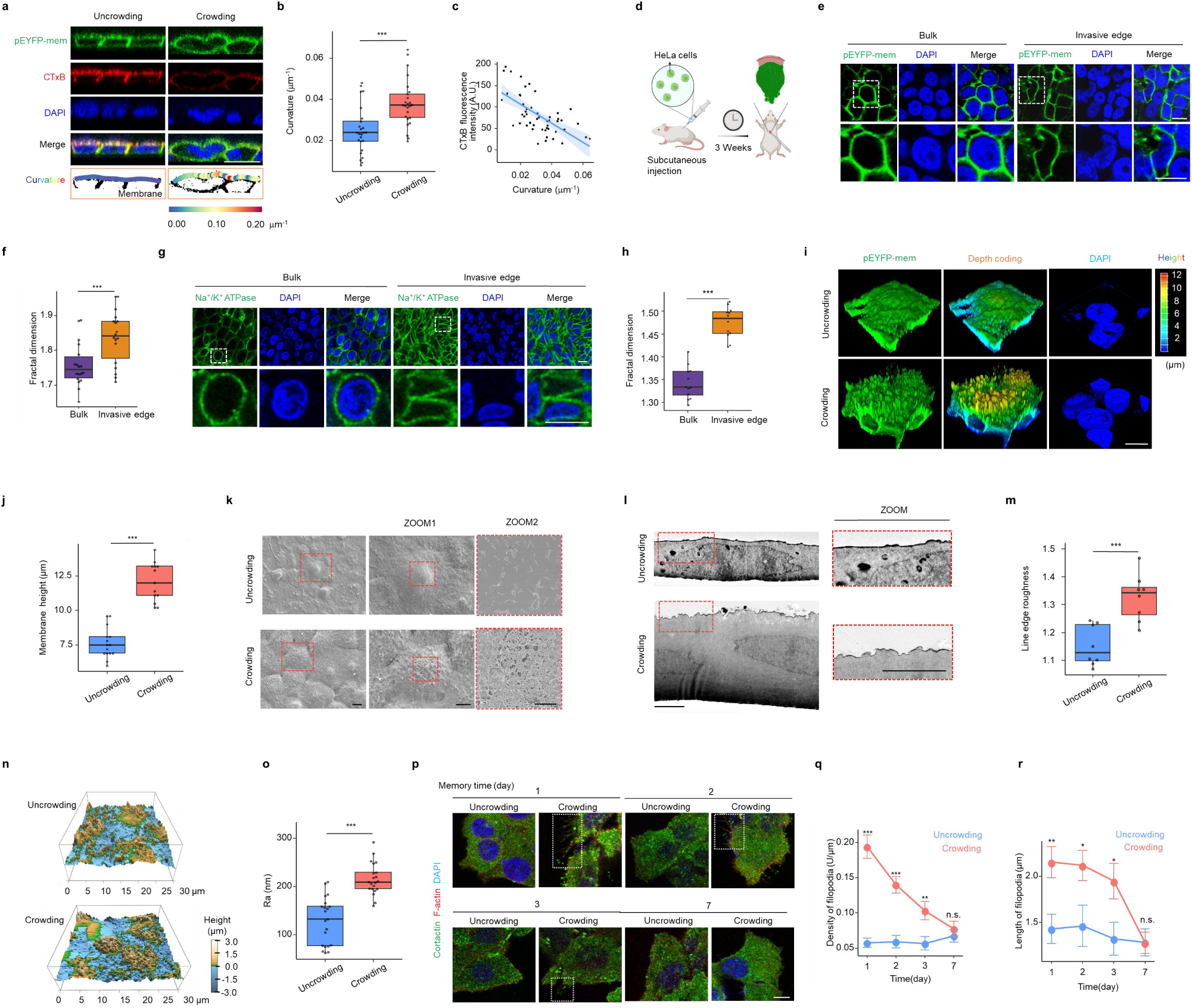
Prolonged crowding triggers a nSCTT of plasma membranes. (**a**) Reconstructed XZ confocal images showing CTxB level and membrane curvature using ImageJ and plugin Kappa in uncrowded and crowded HeLa cells expressing pEYFP-mem. Scale bar: 20 μm. (**b**) Quantitation of membrane curvature in uncrowded and crowded HeLa cells in panel (a). (**c**) Quantitation of correlation between CTxB fluorescence intensity and membrane curvature in panel (a). (**d**) Schematic diagram of subcutaneous nude mouse xenograft model using EYFP-mem+ HeLa cells. After 3 weeks of tumor growth, mouse tissues were harvested and tumor architectures were analyzed for the indicated parameters. (**e**) Representative XY slice image of pEYFP-mem from bulk and invasive edge of nude mouse xenografts. Scale bar: 10 μm. (**f**) Quantitation of correlation between fractal dimension per cell from bulk and invasive edge of nude mouse xenografts. (**g**) Representative XY slice image of sodium potassium ATPase in the uncrowded and crowded regions of human skin cancer tissues. Scale bar: 10 μm. (**h**) Quantitation of correlation between fractal dimension and crowding strain per cell in human skin cancer tissues. (**i**) Confocal 3D reconstruction of the apical membrane of uncrowded and crowded HeLa cells expressing pEYFP-mem. Scale bar: 10 μm. (**j**) Quantitation of membrane height in uncrowded and crowded HeLa cells. (**k**) Scanning electron microscopy (SEM) images of plasma membranes in uncrowded and crowded HeLa cells. Scale bar: 10 and 5 μ m. (**l**) Transmission Electron Microscope (TEM) images of apical plasma membranes revealing the presence of membrane bending in the range of 100 – 500 nm. Scale bar: 5 μm. (**m**) Quantification of line edge roughness in uncrowded and crowded HeLa cells (l). (**n**) The topographic images of HeLa cells by AFM showing the height distribution of the membrane with the size of 30 μm × 30 μm. For uncrowded cells, the surface of membrane was relatively flat. More protruding structures were observed in crowded cells. (**o**) Average roughness (Ra) of the apical plasma membrane was analyzed from 3×3 μm frame ultrastructure images by AFM. (**p**) Representative XY slice image of cortactin distribution in HeLa cells after uncrowding/crowding culture for 5 days and sparse culture for 1,2,3,7 days. Scale bar: 10 μm. (**q, r**) Quantification of filopodia density and length per cell with corresponding memory time in panel (p). Data are presented as mean±SEM; *p < 0.05, **p < 0.01, ***p < 0.005; two-tailed unpaired t-test.

To further assess the effects of prolonged crowding on membrane deformation, we analyzed the surface topography of plasma membranes in EYFP-mem^+^ HeLa cells subjected to prolonged crowding. 3D confocal imaging revealed that the apical plasma membrane of crowded cells exhibited enhanced nanoscale protrusions, in contrast to the much smoother topography observed in uncrowded cells (Fig. 3i, j). This indicates a nanoscale topography transition of plasma membranes from a smooth to a corrugated state, induced by prolonged crowding. To further characterize this transition, we employed scanning electron microscopy (SEM), transmission electron microscopy (TEM), and atomic force microscopy (AFM). SEM and TEM imaging confirmed an enrichment of nanoscale protrusions at the apical plasma membrane in crowded cells compared to uncrowded cells (Fig. 3k-m). AFM analysis demonstrated that the average roughness (Ra), root mean square roughness (Rq), and cellular height were all significantly increased in crowded cells (Fig. 3n, o and Fig. S7a, b). Notably, consistent with the memory retention of invasiveness in cells primed by prolonged crowding, cells primed in crowding culture for 5 days retained elevated cell protrusions (*e.g.*, filopodia) even after 3 days of sparse culture, compared to uncrowded cells (Fig. 3p-r). Taken together, these results demonstrate that prolonged crowding triggers a nanoscale smooth-corrugated topography transition (nSCTT) in plasma membranes, providing a mechanistic basis for the disaggregation of membrane domains and invasion initiation.

### Prolonged crowding increases Laplace pressure to disrupt the aggregation of membrane domains

To investigate the mechanism underlying the nSCTT of plasma membranes triggered by prolonged crowding, we next measured the tension of plasma membranes, which is known to remodel membrane topography by controlling the assembly of curvature-generating proteins^40^. By analyzing the lifetime of a live-cell fluorescent membrane tension probe, Flipper-TR^41^, we found that the tension of the apical plasma membrane in cells exposed to prolonged crowding was significantly reduced compared to that in uncrowded cells (Fig. 4a, b). Plasma membrane tension is known to depend on the contractility of cortical actin^42^. By fluorescence resonance energy transfer (FRET)-based biosensors^43^, we further analyzed the activity of Rho GTPases, which are master regulators of actomyosin structure and dynamics^44^. We observed that the activity of RhoA at the apical side was significantly decreased in crowded cells, while the activity of Rac1 was significantly increased (Fig. 4c, d). Given that RhoA promotes the assembly of stress fibers^45^ and Rac1 mediates branched actin polymerization^46^, these results suggest that prolonged crowding affects membrane tension and cytoskeleton assembly.

**Figure 4.**
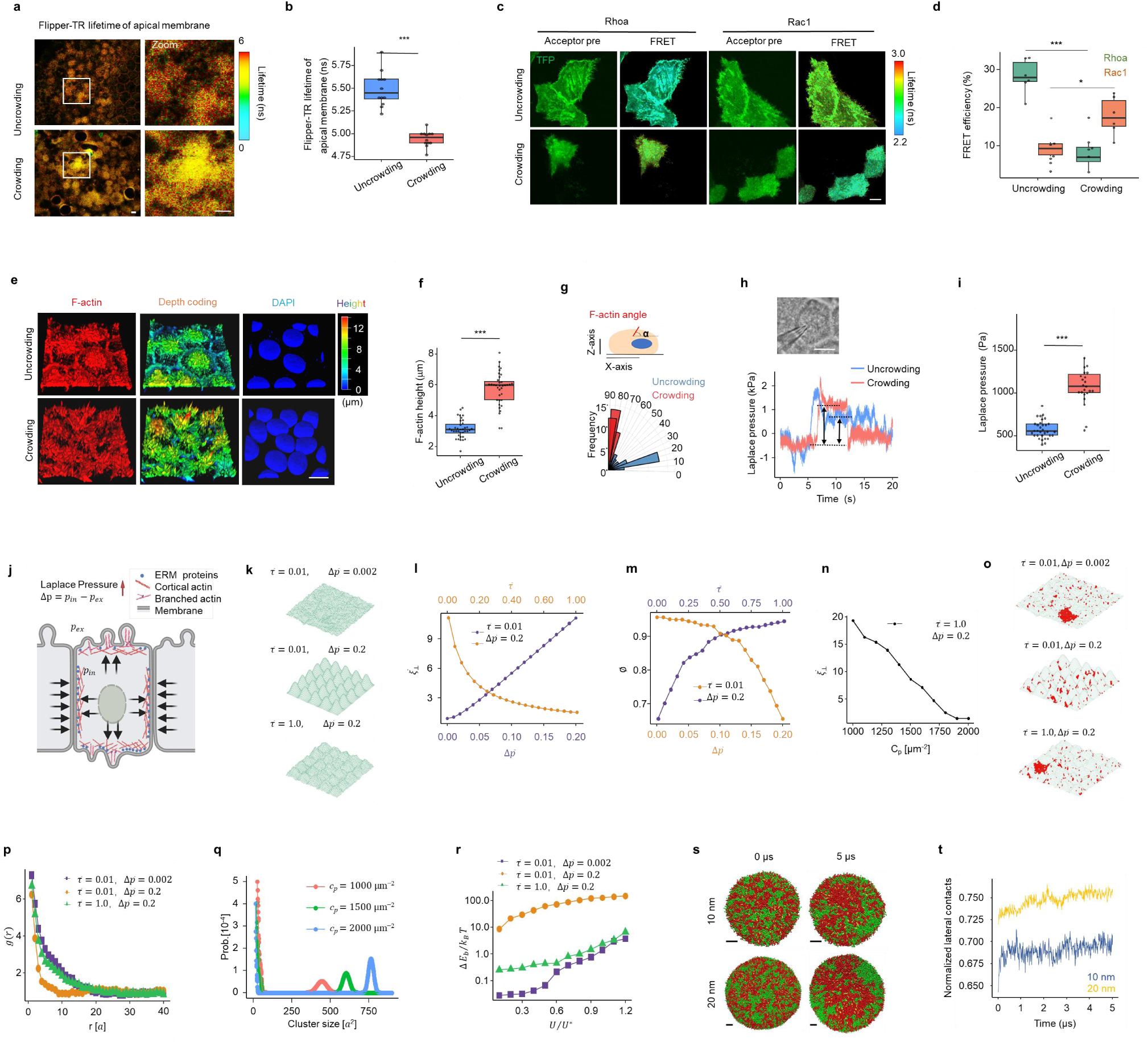
Prolonged crowding triggers nSCTT and disrupts membrane domains by increasing Laplace pressure. (**a**) Representative FLIM images of Flipper-TR lifetime values to analyze membrane tension in uncrowded and crowded cells. Color scale from 0 to 6 ns. Scale bars: 10 µm. (**b**) Quantification of the average lifetime of Flipper-TR from full images. Data are presented as boxplot. (**c**) FRET analysis of uncrowded and crowded HeLa cells expressing the RhoA or Rac1 biosensor. Representative confocal images of over-expressed cells before photobleaching (acceptor pre) are shown. Representative FLIM images of photobleaching in uncrowded and crowded cells are shown. Color scale represents the range of FRET efficiency. Scale bar: 10 µm. (**d**) Quantitation of the fluorescence increase (% FRET efficiency) of RhoA and Rac1 upon photobleaching in uncrowded and crowded cells. Data are presented as boxplot. (**e**) Confocal 3D reconstruction of the apical side of uncrowded and crowded HeLa cells stained for F-actin. Scale bar: 10 µm. (**f**) Quantification of the average height of F-actin from the apical side in uncrowded and crowded cells. (**g**) Actin angle distribution data for cells experiencing uncrowding and crowding displayed as a rose plot in which the absolute value of the angles was compared. (**h**) Quantifying Laplace pressure in uncrowded and crowded HeLa cells using a micro-pressure probe. The baseline reading is close to zero when the microelectrode is not in contact with the cells. A transient pressure spike is recorded as the tip of the microelectrode penetrates the membrane. This is followed by a stable reading for about 5-10 s, before some potential leaks occur, which may cause a gradual drop in the reading with time. Scale bar: 10 µm. (**i**) Quantification of the Laplace pressure of cells experiencing uncrowding and crowding. (**j**) Schematic diagram representing how the increased Laplace pressure controls the protrusions of plasma membranes due to the cellular squeeze of each other. (**k**) Snapshot from Monte Carlo simulations of nanoscale topography of plasma membranes with different membrane tension *τ̅* =*τa*^2^ _*κ*_ and Laplace pressure Δ*p̅* = Δ*pa*^3^ (*κk T*)^−1/2^. (**l**) The rescaled roughness B *ξ̅* =*ξ* (*κ k T*)^1^/ ^2*a*^ as a function of membrane tension and Laplace pressure. (**m**) The ratio of linker ⊥ ⊥ B proteins*ϕ*that bound to the cortical actin as a function of Laplace pressure and membrane tension. (**n**) The rescaled roughness as a function of the density of linker proteins C_p_ that bound to cortical actin. (**o**) Snapshot from Monte Carlo simulations for aggregation of membrane domains for different membrane tension and Laplace pressure as indicated in each figure. (**p**) Pair distribution function g (r) of membrane domains as a function of the distance r for different membrane tension and Laplace pressure as indicated in this figure. (**q**) The probability of cluster size for membrane domains with different density of linker proteins C_p_ that bound to cortical actin. (**r**) The changes of bending energy as a function of the raft-raft contact energy. (**s**) Snapshots of DPPC (red) / DUPC (green) / CHOL (yellow) liposomes with different inner radius at 0 and 5 μs. Scale bar: 4 nm. (**t**) Time evolution of normalized lateral contacts of DUPC lipids for different systems. Data are presented as *p < 0.05, ***p < 0.005; two-tailed unpaired t-test.

To evaluate whether the organization of cortical actin was regulated by prolonged crowding, we performed 3D confocal imaging of cortical actin. We found that the height of apical F-actin was significantly increased in crowded cells compared to uncrowded cells (Fig. 4e, f and Fig. S7c, d). Furthermore, the orientation of actin fibers was altered by prolonged crowding, with fibers in crowded cells exhibiting a more vertical orientation (> 70°related to the base plane), whereas fibers in uncrowded cells were primarily parallel (< 20°) (Fig. 4g). These results suggest that prolonged crowding triggers the remodeling of both plasma membrane and cortical actin.

Previous studies have indicated that membrane-to-cortex attachment (MCA) and cell protrusions (such as blebs) are regulated by the Laplace pressure, the pressure difference across plasma membranes^47^. As a key regulator of cell shape and volume, Laplace pressure influences various cellular processes, including cell migration, proliferation, necrosis, apoptosis, material transportation, and signal transduction^47^. To investigate the biophysical mechanism underlying the nSCTT of plasma membranes induced by prolonged crowding, we analyzed Laplace pressure in cells exposed to different crowding strains using a previously reported micro pressure system^48^. We found that Laplace pressure in crowded cells was significantly higher than in uncrowded cells (Fig. 4h and i).

To further explore the potential impacts of Laplace pressure and membrane tension on the nanoscale topography of plasma membranes, we performed Monte Carlo (MC) simulations of a membrane-cortical actin system (Fig. 4j and Fig. S8). In this MC model, plasma membranes are represented as a fluctuating elastic surface, while cortical actin is modeled as a uniform square mesh framework. The plasma membranes are discretized into square lattices, with each lattice capable of accommodating a transmembrane protein. These transmembrane proteins (linker proteins) connect the plasma membranes to the cortical actin via a harmonic potential^49,50^. The resulting snapshots, as shown in Fig. 4k, clearly demonstrate that increased Laplace pressure ^Δ^ *^p̅^* and decreased membrane tension *^τ̅^* contribute to pronounced bulges and large deformations of the plasma membrane. To quantify these observations, we analyzed the relative roughness *^ξ^*^⊥^ of the cell membrane as a function of both Laplace pressure and membrane tension. The rescaled roughness *^ξ̅^*^⊥^ increased with higher Laplace pressure and lower membrane tension (Fig. 4l). These modeling results, in conjunction with our cellular crowding experiments, provide compelling evidence that the nanoscale topography of plasma membranes is precisely regulated by the interplay between Laplace pressure and membrane tension.

In our MC model, we observed that the detachment of membrane from the cortical actin is regulated in response to changes in Laplace pressure ^Δ^ *^p̅^* and membrane tension *^τ̅^*. Specifically, at a fixed value of *^τ̅^*, when the applied ^Δ^*^p^* exceeds a critical threshold, the proportion *^ϕ^* of linker proteins bound to the cortical actin abruptly drops to zero, indicating complete detachment of the cell membrane from the cortical actin. A similar behavior is observed when *^τ̅^* is reduced to a critical value at fixed ^Δ^*^p̅^* (Fig. 4m). Additionally, increasing the density of linker proteins (C_p_) significantly reduces membrane roughness under constant ^Δ^ *^p̅^* and *^τ^* (Fig. 4n), underscoring the essential role of MCA in the regulation of the nanoscale topography of plasma membranes through the interplay of Laplace pressure and membrane tension. In summary, these findings suggest that the nSCTT of plasma membranes arises from the membrane-to-cortex detachment, driven by an increase in Laplace pressure and a decrease in membrane tension. This mechanism highlights the critical influence of mechanical forces and molecular interactions in shaping membrane morphology.

To further explore the regulation of membrane domain aggregation by Laplace pressure and membrane tension, we incorporated membrane domains into our computational model. These domains exhibit short-range attractive interactions and undergo dynamic fission and merging processes. The distribution of membrane domains is significantly influenced by changes in Laplace pressure ^Δ^ *^p̅^* and membrane tension *^τ^*. At small ^Δ^ *^p̅^*, membrane domains tend to coalesce into a single large domain. However, as Laplace pressure increases, this single domain separates into multiple smaller domains (Fig. 4o). To quantitatively analyze the spatial distribution of membrane domains, we calculated the pair distribution function *g*(r), where a higher *g*(r) value indicates a greater probability of finding two membrane domains at a distance r. As shown in Fig. 4p, an increase in ^Δ^ *^p̅^* results in a larger initial spike in g (r) (where (g(r)>1), suggesting that elevated Laplace pressure reduces the propensity for membrane domain aggregation. Conversely, an increase in membrane tension also leads to a larger initial spike of g(r), indicating that higher membrane tension promotes the aggregation of membrane domains (Fig. 4p). Next, we explored the influence of linker proteins on membrane domain aggregation within our MC simulations. Our analysis indicated that augmenting the density of linker protein C_p_ enhances the aggregation of membrane domains (Fig. 4q). These results underscore that the aggregation of membrane domains is disrupted by increased Laplace pressure and membrane tension under conditions of prolonged crowding.

To elucidate the mechanism governing the regulation of lipid membrane domain aggregation by Laplace pressure and membrane tension, we estimated the bending energy of the multicomponent membrane using a discretized Helfrich Hamiltonian^51^. At low raft-raft contact energy *U*, the membrane domains are uniformly distributed across the cell membrane. As *U* increases, these domains tend to coalesce into a single, larger domain. The change in bending energy Δ*E*_b_ rises with the increasing raft-raft contact energy *U*, suggesting that the aggregation of membrane domains is energetically unfavorable (Fig. 4r). This analysis of bending energy reveals that both elevated Laplace pressure and reduced membrane tension increase the energy change Δ*E*_b_, thereby raising the energetic cost for membrane domain aggregation.

To further investigate the effects of membrane curvature on the aggregation of membrane domains, we conducted μs-scale coarse-grained (CG) molecular dynamics (MD) simulations of three-component vesicles (DPPC: 50%; DUPC: 30%; CHOL: 20%) with two different radii (10 nm and 20 nm) using the Martini CG model^52,53^. As illustrated in the system snapshots and normalized lateral contacts of DUPC lipids (Fig. 4s and t), DPPC/DUPC/CHOL vesicles underwent pronounced aggregation of membrane domains in both vesicle sizes. However, the larger vesicles (r = 20 nm), characterized by reduced membrane curvature, displayed significantly enhanced aggregation of membrane domains. These results suggest that the local nanoscale topography with higher membrane curvature inhibits the aggregation of membrane domains. Collectively, these findings highlight that the aggregation of membrane domains is disrupted under prolonged crowding condition through a complex interplay of mechanical forces and molecular interactions.

### Suppressing the pressure-sensation of membrane domains inhibits tumor invasion

To empirically validate the predictions from our simulations, we investigated the distribution of ERM proteins (ezrin, radixin, and moesin), which are known to mediate MCA by linking plasma membranes to cortical actin^54,55^, in the cellular crowding model. Immunofluorescence staining with an antibody specific to phosphorylated ERM (pERM), the active membrane- and actin-bound form of ERM, revealed a progressive reduction in pERM levels at the apical membrane as crowding strain increased (Fig. 5a and b). Furthermore, this reduction in apical pERM levels in crowded cells persisted for 3 days after transitioning to sparse culture conditions (Fig. 5c and d). These findings demonstrate that prolonged crowding leads to a sustained decrease in activated ERM levels, indicating a form of mechanomemory in cancer cells.

**Figure 5.**
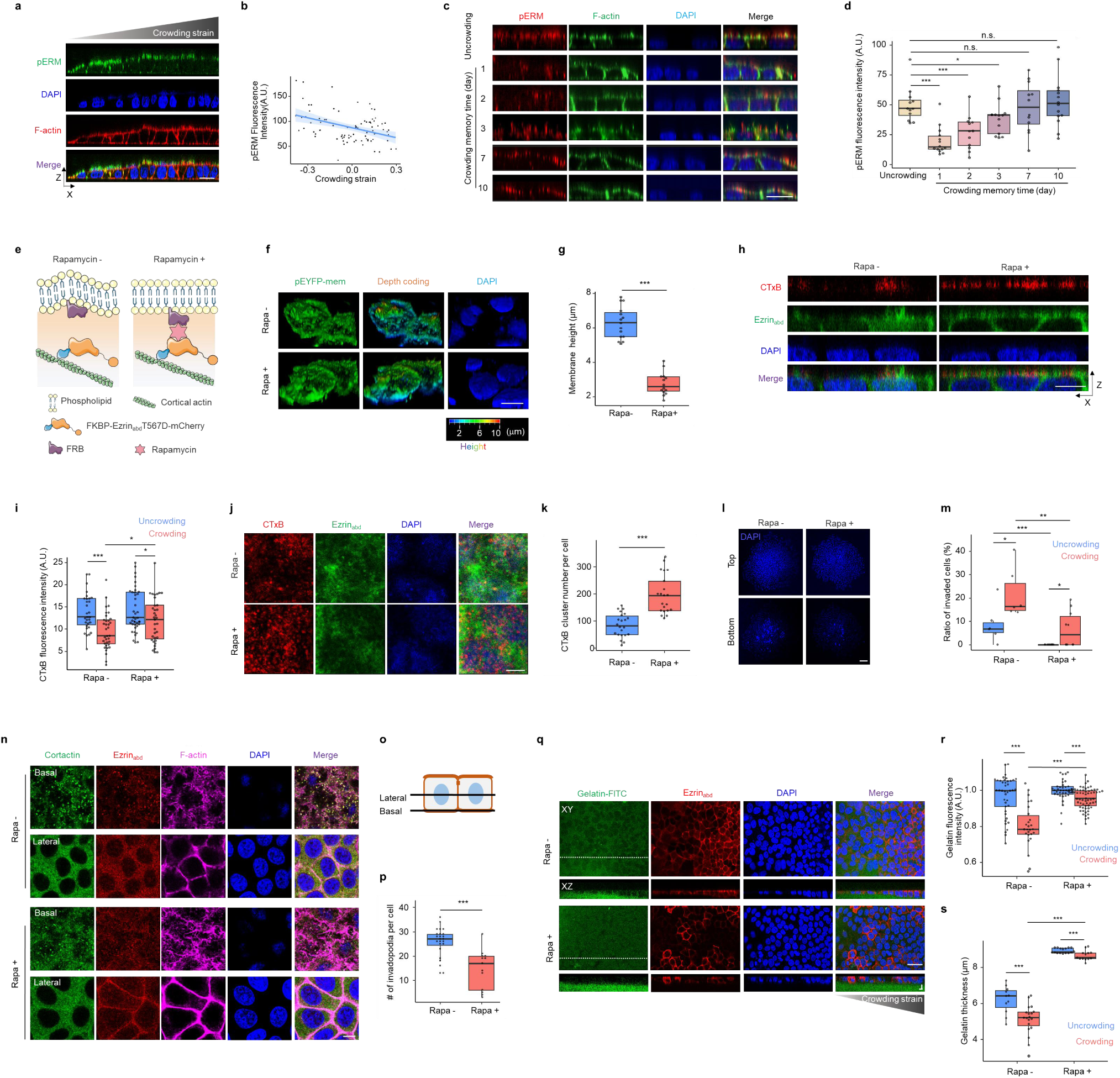
Suppressing the pressure-sensation of membrane domains inhibits crowding-initiated cancer cell invasiveness. (**a**) Reconstructed XZ confocal images of pERM localization at the apical side in uncrowded and crowded HeLa cells. Scale bar: 20 μm. (**b**) Quantification of pERM fluorescence intensity per cell with corresponding crowding strain. (**c**) Reconstructed XZ confocal images of pERM localization at the apical side of HeLa cells after uncrowding/crowding culture for 5 days and sparse culture for 1, 2, 3, 7, 10 days. Scale bar: 20 μm. (**d**) Quantification of pERM fluorescence intensity with corresponding memory time from uncrowded and crowded groups. (**e**) Schematic diagram for a FKBP-EzrinabdT567D construct to acutely increase cortical actin by recruitment of the EzrinabdT567D to the plasma membranes after rapamycin treatment. (**f**) Confocal 3D reconstruction of the apical membrane of FKBP-EzrinabdT567D-mCherry+; EYFP-mem+ HeLa cells treated with rapamycin or vehicle. Scale bar: 10 µm. (**g**) Quantification of the average height of membrane from the apical side in control and rapamycin treated cells. (**h**) Reconstructed XZ confocal images showing CTxB fluorescence intensity in FKBP-EzrinabdT567D-mCherry+ HeLa cell sheets treated with rapamycin or vehicle. Scale bar: 10 μm. (**i**) Quantitation of CTxB fluorescence intensity in uncrowded and crowded cells treated with rapamycin or vehicle. (**j**) Representative XY confocal images of CTxB staining at the apical membrane of FKBP-EzrinabdT567D-mCherry+ HeLa cell sheets treated with rapamycin or vehicle. Scale bar: 5 μm. (**k**) Quantitation of the number of CTxB cluster per cell treated with rapamycin or vehicle. (**l**) Transwell matrigel invasion assay of FKBP-EzrinabdT567D-mCherry+ HeLa cell sheets after treatment with rapamycin or vehicle. Representative DAPI images of cells that accumulated on the top (uninvaded) and bottom (invaded) surface of the insert membranes. Scale bar: 100 μm. (**m**) Ratio of invaded cells in FKBP-EzrinabdT567D-mCherry+ HeLa cell sheets treated with rapamycin or vehicle in panel (l). (**n**) Representative XY slice image of cortactin and F-actin staining in FKBP-Ezrinabd T567D-mCherry+ HeLa cell sheets treated with rapamycin or vehicle. Scale bar: 10 μm. (**o**) Schematic diagram of lateral and basal cross-sections in HeLa cell sheets. (**p**) Quantification of invadopodia density per cell in control and rapamycin groups in panel (n). (**q**) FKBP-EzrinabdT567D-mCherry**+** HeLa cells were plated on FITC-conjugated gelatin hydrogels. Representative XY and XZ slice images of gelatin-FITC and EzrinabdT567D-mCherry in cell sheets after rapamycin treatment. Scale bar: 10 µm. (**r, s**) Quantification of Cy5-conjugated gelatin fluorescence intensity and thickness in uncrowded and crowded cells treated with rapamycin or vehicle in panel (q). Data are presented as *p < 0.05, **p < 0.01, ***p < 0.005; two-tailed unpaired t-test.

Our simulation results revealed that enhanced MCA confers resistance against the nSCTT of plasma membranes and prevents the disaggregation of membrane domains under prolonged crowding conditions (Fig. 4n and q). To experimentally validate this prediction, we employed a Lyn-FRB and Ezrin_abd_-FKBP activation system^42^, which incorporates a constitutively active F-actin binding domain of Ezrin (Ezrin_abd_T567D). Following rapamycin treatment, Ezrin_abd_T567D was efficiently recruited to the membrane, leading to a significant increase in cortical actin density (Fig. 5e, and Fig. S9a-c). Acute reinforcement of MCA through recruitment of FKBP-Ezrin_abd_T567D to plasma membranes resulted in a marked reduction in both membrane protrusions and apical F-actin height in crowded cells (Fig. 5f-g, and Fig. S9d-e). Furthermore, enhanced MCA via FKBP-Ezrin_abd_T567D activation significantly increased the aggregation of membrane domains, as evidenced by both fluorescence intensity and the number of CTxB clusters in crowded cells treated with rapamycin compared to untreated cells (Fig. 5h-k). These results demonstrate that the aggregation of membrane domains, disrupted by prolonged crowding, can be rescued by inhibiting the nSCTT of plasma membranes through the enhancement of MCA.

Next, we investigated whether experimentally enhancing MCA could suppress the invasion driven by prolonged crowding in the cellular crowding model. Transwell invasion assays revealed that enhancing MCA through FKBP-Ezrin_abd_T567D activation significantly inhibited the invasion of cancer cells exposed to prolonged crowding (Fig. 5l and m). Immunofluorescence staining of cortactin at the basal membrane demonstrated that the number of invadopodia per cell in crowded cells was markedly reduced by FKBP-Ezrin_abd_T567D activation compared to control cells (Fig. 5n-p). Furthermore, FKBP-Ezrin_abd_T567D activation also suppressed ECM degradation, as evidenced by experiments using Cy5-conjugated gelatin hydrogels (Fig. 5q-s). These results suggest that enhancing membrane domain aggregation by strengthening MCA efficiently rescues the prolonged crowding-driven invasiveness.

To evaluate whether enhancing MCA could suppress tumor invasion *in vivo,* we established a subcutaneous mouse xenograft model using FKBP-mCherry^+^ or FKBP-Ezrin_abd_T567D-mCherry^+^ HeLa cells. At one-week post-cancer cell inoculation, the mice were treated with rapamycin (Fig. 6a). Consistent with the *in vitro* data (Fig. S9a), F-actin was recruited to the cell membrane in FKBP-Ezrin_abd_T567D-mCherry^+^ xenografts after two weeks of rapamycin administration, contrasting with FKBP-mCherry^+^ xenografts (Fig. 6b). Immunofluorescence analysis using anti-sodium potassium ATPase (Na^+^/K^+^ ATPase) antibody for membrane labeling revealed a significant reduction in the fractal dimension of plasma membranes in crowded cells of FKBP-Ezrin_abd_T567D-mCherry^+^xenografts compared to FKBP-mCherry^+^ xenografts (Fig. 6c). Furthermore, elevated levels of membrane domain markers (Caveolin-1 and Flotillin-1) were detected in FKBP-Ezrin_abd_T567D-mCherry^+^ cells compared to FKBP-mCherry^+^ cells (Fig. 6d-g). These findings demonstrate that enhanced MCA effectively suppresses the nSCTT of plasma membranes and mitigates membrane domain disaggregation in crowded cancer cells within xenografts.

**Figure 6.**
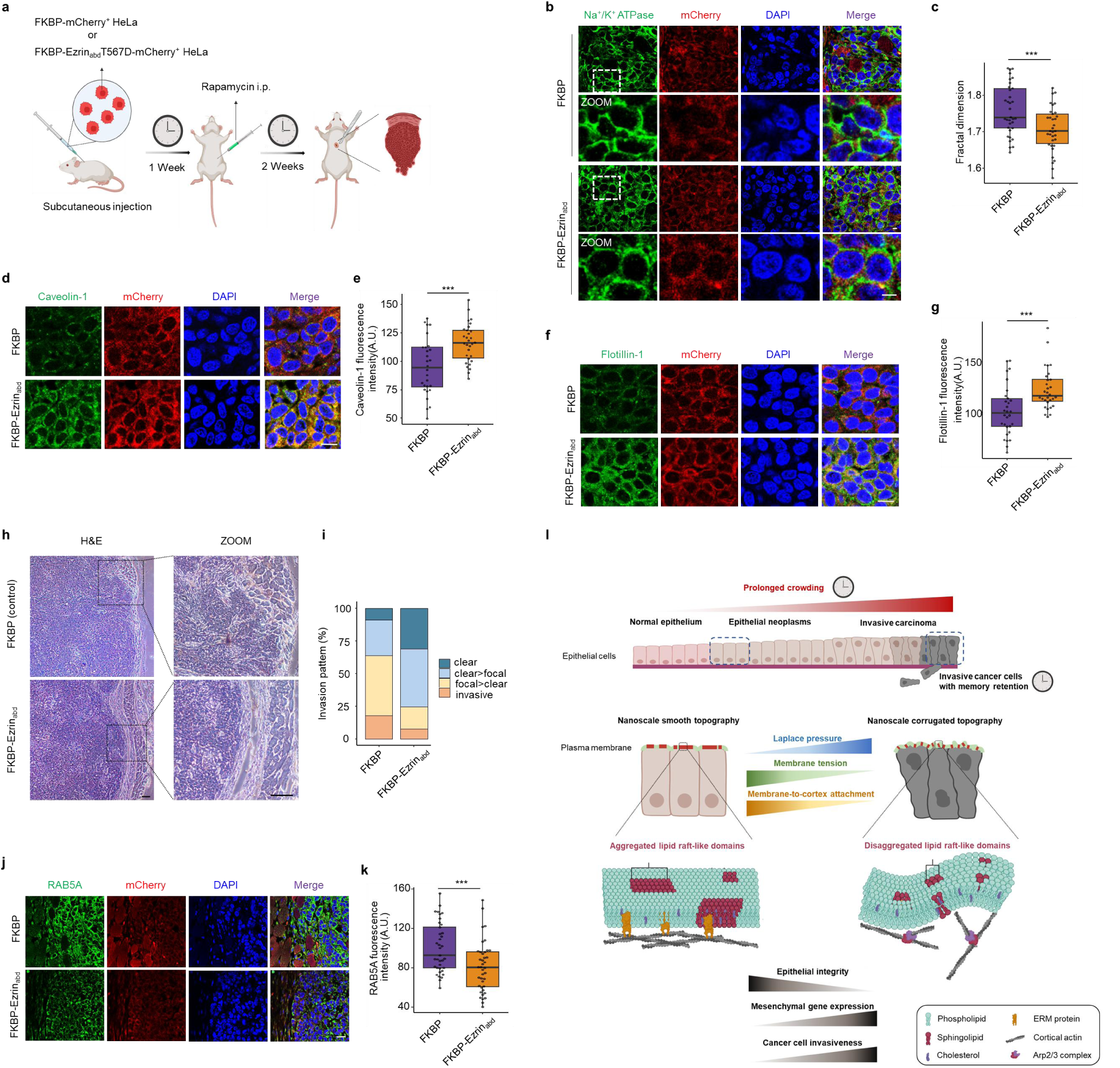
Suppressing the pressure-sensation of membrane domains inhibits tumor invasion in a mouse xenograft model. (**a**) Schematic diagram of subcutaneous nude mouse xenograft model using FKBP-Ezrin_abd_T567D and FKBP (Control) over-expressed HeLa cells. After 1 week of tumor growth, mice were treated with rapamycin via i.p. three times per week. Tissues were harvested and tumor architectures were analyzed for the indicated parameters. (**b**) Representative XY slice images of sodium potassium ATPase and mCherry in mouse xenograft treated with rapamycin. Scale bar: 5 μm. (**c**) Quantitation of fractal dimension of HeLa cells expressed FKBP-Ezrin_abd_T567D and FKBP in mouse xenografts. (**d**) Immunofluorescence analysis of Caveolin-1 in mouse xenografts treated with rapamycin. Scale bar: 10 μm. (**e**) Quantification of Caveolin-1 fluorescence intensity of tumor cells in panel (d). (**f**) Immunofluorescence analysis of Flotillin-1 in mouse xenografts treated with rapamycin. Scale bar: 10 μm. (**g**) Quantification of Flotillin-1 fluorescence intensity of tumor cells in panel (f). (**h**) Representative images of HE staining in mouse xenografts treated with rapamycin. scale bar: 100 µm. (**i**) Invasion pattern of mouse xenografts in panel (h). Invasion was classified as “clear” (distinct border between muscle and tumor), “clear>focal” (more clear borders than areas with focal invasions), “focal>clear” (more focal invasions than clear borders), or “invasive” (no clear borders). Percentages of each category are given. Only samples with sufficient surrounding muscle tissue were evaluated. (**j**) Immunofluorescence analysis of RAB5A and mCherry in mouse xenografts treated with rapamycin, scale bar: 20 µm. (**k**) Quantification of RAB5A fluorescence intensity in the invasive edge of tumor in mouse xenografts. (**l**) Schematic diagram of pressure-sensing membrane domains triggered by prolonged crowding driving cancer cell invasiveness. The supported membrane consists of non-membrane domains containing unsaturated phospholipids and lipid membrane domains by association of sphingolipid and cholesterol molecules. Under uncrowded conditions, aggregated membrane domains reside in nanoscale smooth topography, maintaining a flat plasma membrane sustained by membrane-to-cortex attachment. In contrast, under prolonged crowding conditions, membrane domains become disaggregated are confined to regions with nanoscale corrugated topography with plasma membrane protrusions. This transition in membrane domain organization and topography is critical for driving cancer cell invasiveness. Schematic images (a, l) were created with BioRender.com Data are presented as ***p < 0.005; two-tailed unpaired t-test.

Next, we investigated the invasive growth patterns in xenografts with or without strengthening MCA. Hematoxylin and eosin (H&E) staining demonstrated that FKBP-mCherry^+^ xenografts exhibited deeper invasion into the muscle layer (Fig. 6h and i). In contrast, FKBP-Ezrin_abd_T567D-mCherry^+^ xenografts displayed well-defined pushing tumor borders, effectively segregating cancer cells from the adjacent muscle layer with only minor focal invasions (Fig. 6h). Immunofluorescence staining for RAB5A further reveled that the invasive edge of tumors in FKBP-Ezrin_abd_T567D-mCherry^+^ xenografts exhibited reduced invasiveness compared to FKBP-mCherry^+^ xenografts (Fig. 6j and k). Additionally, the inhibitory effect of MCA enhancement on tumor invasion was confirmed by comparing FKBP-Ezrin_abd_T567D-mCherry^+^ HeLa cell xenografts with and without rapamycin treatment (Fig. S10). Taken together, these results demonstrate that preventing membrane domain disaggregation through MCA enhancement effectively suppresses cancer cell invasiveness *in vivo*.

## DISCUSSION

Growing in confined spaces, cancer cells push and stretch solid components of the surrounding tissue and thus experience situations of crowding^8^. Recent studies have shown that crowding triggers cell extrusion and induces cell migration during embryonic development and tumor progression^56–59^. Here, using a spontaneous crowding model composed of a freely growing monoclonal cell sheet, we found that prolonged crowding drives cancer cell invasiveness with memory retention by triggering a nanoscale topography transition of plasma membranes from a smooth to a corrugated state. Combining mechanical modeling, genetic manipulations, and biophysical measurements, we further found that the prolonged crowding-triggered nSCTT of plasma membranes disrupts the aggregation of membrane domains to drive tumor invasion (Fig. 6l).

Plasma membranes of cancer cells exhibit complex and irregular shapes. Analyses using fractal geometry are already used to efficiently estimate the geometrical shapes observed during tumor progression and for ascertaining correlations with pathological processes^60^. For example, skin cancer can be modeled by calculating fractals to evaluate the invasiveness of cancer cells^60^. The fractal dimension of AFM maps is analyzed for three stages of progression towards cervical cancer, from normal through immortal to malignant cells^61^. Here, we identified that prolonged crowding increases the Laplace pressure and decreases membrane tension, which induces nanoscale protrusions at the apical plasma membrane of cancer cells. The aggregation of membrane domains suppressed by the nSCTT of plasma membranes disrupts epithelial integrity and drives cancer cell invasiveness.

Although controversies about the composition, properties, and even the existence of membrane domains remain unresolved^62^, previous studies have demonstrated that most transmembrane proteins are found in membrane domains to serve as signaling platforms or connect cortical actin for controlling cell behaviors^63^. Recent observations of model membranes reported that macroscopic membrane domains exist in membrane regions with nanoscale smooth topography^39^. Another study has shown that pharmacological disruption of rafts using MβCD decreases CD44 retention inside membrane domains and promotes CD44 interaction with ezrin to drive cell migration^64^. In addition, oncogenic mutants of RAS proteins (encoded by *HRAS, KRAS, NRAS*) are localized to the inner leaflet of the plasma membranes, and the assembly of RAS nanoscale clusters is known to depend on the interactions between plasma membrane lipids and RAS molecules^65,66^. In light of our findings, RAS activation may be driven by the aggregation of membrane domains in cells experiencing the nSCTT of plasma membranes triggered by prolonged crowding.

As a transmembrane pressure of cells, the Laplace pressure which is the difference between intracellular and extracellular pressures can rapidly reprogram cell shape and regulate cell migration^47,67^. Membrane tension, which arises from the combined contributions of osmotic pressure, in-plane tension, and cytoskeletal forces, remodel membrane topology and influence cellular function^40^. In tumor environments, the mechanical characteristics of crowding lead to changes in both the Laplace pressure and membrane tension of cancer cells. These cells’ behavior regulation in crowded environments maybe mediated by mechanosensitive ion transporters and channels affected by Laplace pressure and membrane tension^68,69^. Our findings demonstrate that the nanoscale smooth-corrugated topography transition of plasma membranes induced by crowding can significantly enhance cancer cell invasiveness.

Our experiments demonstrated that plasma membranes exhibit a configuration of nanoscale topography that is consistent with cortical actin protrusion. Recent studies have reported that tumor cells migrate through crowded environments via large bleb protrusion controlled by actin filament at the cell front to break apart ECM^70,71^. ERM proteins, which tether the membrane to cortical actin of cells, restrict local membrane protrusions and inhibit cancer cell migration^42^. This is consistent with our experimental findings showing that enhancing membrane-to-cortex attachment by FKBP-Ezrin_abd_T567D significantly inhibits the prolonged crowding-induced nSCTT of plasma membranes and the invasiveness of cancer cells.

The geometrical irregularity of tumor boundaries has been implicated in the development and progression of cancers such as skin, breast, and lung cancer^60,72,73^. The application of fractal geometry for analyzing the surface of human cervical and breast cancer cells has shown considerable promise for estimating cancer stages^61,74,75^. Abnormal changes in the shape of the plasma membranes, including bending and protrusion, promote the migration and invasion of cancer cells^54^. Our experiments and modeling results demonstrate that prolonged crowding drives cancer cell invasiveness by triggering a nSCTT of plasma membranes. Inhibiting the nSCTT by enhancing MCA effectively suppresses cancer cell invasiveness induced by prolonged crowding. Our findings provide clues for potential clinical strategies targeting the nanoscale topography of plasma membranes for preventing carcinoma invasion. In particular, given the essential roles of ERM proteins in regulating crowding-driven invasiveness, it will be interesting to further explore if certain small molecules or gene-targeting entities could be identified to suppress ERM dephosphorylation and thus reinforce MCA when delivered to the tumor tissues. These new agents may hinder the progression of tumors toward a more malignant stage by normalizing the plasma membrane topography of cancer cells subjected to prolonged crowding.

## METHODS

### Mouse lines

SPF male BALB/c nude mice aged 3-4 weeks and weighing 16-18 g were purchased from Vital River Laboratory Animal Technology Co., Ltd. Mice were bred and reared in the animal facility of Tsinghua University at 22 °C with a 12-hour light/dark cycle (lighting time 7:00-19:00). Food and water are freely available. All animal studies were conducted under the guidance of the Animal Care and Utilization Committee (IACUC) of Tsinghua University. According to the National Institutes of Health “Animal Ethical Use Guidelines”, the experimental procedure has been approved by the Laboratory Animal Care and Use Management Committee of Tsinghua University and the Beijing Municipal Science and Technology Commission (SYXK-2019-0044).

### Human tissues

Samples of human breast cancer, skin cancer, and cervical cancer (paraffin sections) were obtained from patients who had undergone surgery at the Dezhou Second People’s Hospital. The cases were classified according to the World Health Organization classification criteria of the tumors. The samples were collected with patient consent, following approval by the Institutional Committee for the Welfare of Human Subjects.

### Maintenance of cell lines

HeLa, A431, and MC38 cells were cultured in Dulbecco’s modified eagle medium (DMEM) medium (containing 4.5 g/L glucose, L-glutamine and sodium pyruvate) supplemented with 10% fetal bovine serum (FBS; Life technologies, CA, USA), and 100 IU/mg penicillin-streptomycin (Life technologies, CA, USA), and 1% (v/v) non-essential amino acids (NEAA; Life technologies, CA, USA).

### Plasmid construction and transfection

Cells were transiently transfected with plasmid DNA (pEF1-Lifeact-mCherry, pEYFP-membrane, pTriEx4-Rac1-2G and pTriEx-RhoA-2G) using Lipofectamine 2000 (PN: 11668030, Invitrogen), according to the manufacturer’s protocol. pTriEx4-Rac1-2G and pTriEx-RhoA-2G were a gift RhoA-2G-control were constructed by deleting the coding sequence of acceptor Venus. C1-mCitrine-FKBP-EZRabd(t567D), pCAG-Lyn11-FRB and YFP-FKBP were a gift from Tobias Meyer (Addgene plasmid no. 155227, 155228, and 20175). pLV[Exp]-Puro-CMV>mCherry/FKBP-EZRabd(t567d), pLV[Exp]-Puro-CMV>mCherry/FKBP and pLV[Exp]-Puro-EF1A>Lyn-FRB-HA were constructed by VectorBuilder.

### Drug treatments

Pharmacological inhibitors and chemical compounds were used at the following concentrations: 100 μM Cholesterol (Merck, #57-88-5), 100 μM MβCD (membrane domains inhibitor; Merck, #128446-36-6). Rapamycin (MCE, #53123-88-9). Rapamycin was dissolved in DMSO for treatment in cells (5 μM) and dissolved in 5.2% polyethylene glycol and 5.2%Tween-80 for treatment in mice (2.0 mg/kg).

### Subcutaneous mouse xenograft model

The mouse xenograft model was established by subcutaneous inoculation of 5×10^6^ HeLa cells into the suprascapular region of 6-week-old nude mice (10 per group). When tumors reached 100–200 mm^3^ after 1 week, mice were randomly assigned to receive intraperitoneal injection of vehicle (0.25 % polyethylene glycol, 0.25 % tween 80) or rapamycin (2 mg/kg) every other day. Tumors were measured weekly using calipers and volume was calculated as (length×width^2^)/2. Mice were euthanized after 2 weeks of treatment, and tumors were excised and fixed with 4% paraformaldehyde.

### Immunofluorescence and histological analysis

Cells grown on glass bottom or confocal dishes were fixed with 4% paraformaldehyde, permeabilized with 0.1% Triton X-100 for 5 min. Cells were incubated with primary antibodies at the optimal concentrations (according to the manufacturer’s instructions) at 4 °C overnight. Following three washes with PBS, samples were incubated with the appropriate secondary antibodies: 488/568/633 immunoglobulin G (IgG; H+L) and/or Alexa Fluor 568/647 phalloidin (Invitrogen) for 1 h at room temperature. Samples were again washed three times with PBS and mounted with 4’,6-diamidino-2-phenylindole (DAPI, Invitrogen) for 10 min at room temperature. Confocal images were taken on the Leica microscope. Experiments were replicated at least three times. Acquisition of fluorescent images was carried out using a Leica TCS SP8 AOBS Confocal laser-scanning microscope equipped with a 10×, 40×, or 63×objective (Leica, Germany).

Prior to embedding in paraffin, mouse tissues were fixed in 4% paraformaldehyde in PBS and dehydrated. For histological analysis, 6 µm sections were cut and stained with Hematoxylin and Eosin. For immunofluorescence analysis, 6 µm sections were incubated for 20 min in 10 mM sodium citrate buffer, pH 6.0 at 90 °C to retrieve antigens on paraffin-embedded tissue samples. After 1 h incubation in 5% fetal calf serum, sections were incubated overnight with diluted primary antibodies, washed and further incubated for 2 h at room temperature with appropriate secondary antibodies. Nuclei were stained with DAPI for 10 min at room temperature. Confocal images were obtained using Leica microscope equipped with a 10×, 40×, or 63×objective. Experiments were replicated at least three times.

### Cholera toxin subunit B (CTxB) staining

HeLa cells were grown on 35 mm confocal dishes and cultured for 14 days. The cells were then rinsed with cold Hank’s balanced salt solution (HBSS)+0.5% BSA and incubated with 0.5 μg/mL CFTM 488A or 633 conjugated CT-B in cold HBSS+0.5% BSA and incubated at 4°C for 30 minutes in the dark. Cells were washed five times with cold HBSS+0.5% BSA and fixed in 4% paraformaldehyde in PBS for 15 minutes at room temperature. Nuclei were stained with DAPI for 10 min at room temperature.

### Transwell invasion and migration assays

For the Transwell invasion assay, briefly, 5-10 cells in 200 µL 10% FBS DMEM were reseeded into the upper chamber of a 24-well Transwell of 8 μm pore size (Corning Inc., Corning, NY, USA) with coated-Matrigel (BD Bioscience, San Jose, CA, USA), and 600 µL medium with 10% FBS was loaded into the well below. After culture for 14 days, the upper chamber was replaced with serum-free medium and later incubated at 37 °C for 48 h. Transwell insert membranes were fixed with 4% paraformaldehyde and stained with DAPI. For the Transwell migration assay, all procedures were similar but without the incubation of Matrigel. The percentage of migrating and invading cells through the filter was imaged under a Leica SP8 Confocal microscope, and measured using the ImageJ software.

### GelMA degradation assay

Gelatin-Cy5 and Gelatin-FITC methacryloyl (GelMA, EFL-GM-90, China), lithium phenyl-2, 4, 6-trimethylbenzoylphosphinate (LAP), and a blue light source (3 W, 405 nm) were purchased from Engineering for Life, Suzhou, China. GelMA was dissolved in PBS at 30% (w/v) containing 0.25% (w/v) LAP. The mixture was transferred to a glass slide and exposed to blue light irradiation for 90 s to crosslink the GelMA. Then the prepared hydrogels were rinsed with PBS three times, and then a single cell was seeded on hydrogels in 24-well plates and cultured for 14 days. Cells were fixed with 4% paraformaldehyde at room temperature for 15 min and washed three times with PBS. The sample was incubated in 0.1% Triton X-100 for 10 min. Subsequently, the samples were stained for actin with phalloidin for 60 min and stained for nuclei with DAPI for 10 min.

### RNA seq analysis

HeLa cells were primed for 20 days in crowding or uncrowding culture, followed by exposure to sparse culture for different times (0, 1, 3, or 10 days). The samples were collected and sent to the Beijing Genomics Institute (BGI, China) for RNA sequencing performed on the BGISEQ-500 platform. The data was analyzed using Dr. Tom system multi-omics interactive system (BGI, China). Q-value was obtained by false discovery rate (FDR) correction of the P-value. Differentially expressed genes (DEGs) (Q≤0.05, |log2FC|≥1) were analyzed by DEseq2 software. Volcano plot visualization of gene expression patterns was performed using R and results with Q ≤ 0.05 were considered statistically significant.

### Atomic force microscopy (AFM)

HeLa cells were grown on 35 mm confocal dishes and cultured for 14 days. Cell samples were fixed with 4% paraformaldehyde, after washing cells with PBS to remove potential impurities on the cell surface, and then kept in PBS. AFM experiment was performed in PBS buffer solution at room temperature in PeakForce Tapping mode by scanning probe microscope Asylum MFP-3D-SA (Asylum Research, USA). A PeakForce qp-BioAC probe (nominal spring constant 0.06-0.18 N/m, Nanosensors, Neuchatel, Switzerland) was used to image the cell surface. The scanning parameters were as follows: scan size of 30 μm, scan rate of 0.1 Hz, set point of 162.21 mV, integral gain of 611.37, drive amplitude of 2 V. All images were taken at a resolution of 256×256 pixels. The scan area depended on the size of the HeLa cell and ranged from 30×30∼50×50 μm^2^. Image processing and data analysis were performed by the Asylum MFP-3D-SA software.

### Focused ion beam and scanning electron microscopy

The fixation of micropatterned HeLa cells was performed at room temperature for 15 minutes using a 2.5% v/v glutaraldehyde (Electron Microscopy Sciences) solution in PB buffer. After washing the samples three times with PB buffer, the samples were osmicated with 1% osmium tetroxide/1.5% potassium ferricyanide in distilled water for 30 minutes. The samples were then washed three times with distilled water and then dehydrated through a graded ethanol series. After dehydration, samples were infiltrated with Pon 812 Resin (SPI) by incubating the samples in a diluted series of ethanol-Pon 812 at a 1:1, 1:2, and 1:3 ratio for 1 hour for each, followed by overnight in pure resin. The pure resin was changed once in the first hour, then the samples were incubated in an oven at 60 °C for 48 hours. 70 nm sections were cut by ultramicrotome (Leica EM UC7) and stained with uranyl acetate (UA) and lead citrate, then imaged by TEM. To visualize the cell surface, the resin covering the cell was removed using acetone, and the cells filled with resin were polymerized at 60 °C for 48 hours. 10nm gold was coated before imaged by SEM. After that, resin was re-applied, and 70 nm Cross section of the cells were cut for TEM imaging.

### Laplace pressure calculation

Laplace pressure measurements were conducted using the 900A micro pressure system (WPI) based on the servo-null method, following the manufacturer’s instructions. A microelectrode was created from a glass capillary (0.75 mm inner diameter/1.0 mm outer diameter) using a micropipette puller (PC-100, Narishige). The one-stage pull mode was employed with the following settings: Heat 50V, Weights: 250 g. Before the measurement, the microelectrode was calibrated using the calibration chamber and pressure source. The microelectrode was filled with a 1M NaCl solution, while the calibration chamber contained a 0.1M NaCl solution. To perform the measurements, the microelectrode was mounted to a piezo-driven xyz micromanipulator (SN-PCZ-50R, WPI) located in an environmental chamber (37 °C, 5% CO2). A four-channel AD converter was used to record the pressure signal. The microelectrode tip was inserted into the cells at a 45-degree angle and then slightly retracted to release compression on the cells. This position was maintained for at least 10 seconds, and the Laplace pressure was determined as the average pressure during this this period.

### Membrane tension measurements

Cell membrane tension was measured using Flipper-TR fluorescent tension probe (SC020, Cytoskeleton, Inc.). HeLa cells were cultured on the gelatin-coated 35 mm confocal dishes for 14 days until the cell sheet was formed. Cells were then treated with 1 mM Flipper-TR at 37 °C for 15 minutes to achieve appropriate labeling prior to imaging. The fluorescence lifetime of Flipper-TR was measured by using an Olympus fluorescence lifetime imaging microscope (FLIM, FV-1200, Japan). Excitation was performed using a pulsed 488-nm laser operating at 40 MHz, and the emission signal was collected through a 550-650-nm bandpass filter using a HyD SMD detector. Lifetimes of Flipper-TR were extracted from FLIM images using SymPhoTime 64 software (PicoQuant).

### Fluorescence resonance energy transfer (FRET)

HeLa cells were cultured on the gelatin-coated 35 mm confocal dishes for 14 days and transfected with genetically encoded biosensors expressing Rac1-2G-control, Rac1-2G, RhoA-2G, or RhoA-2G-control. After 48 hours, acquisition of fluorescent images and FRET experiments were carried out using an Olympus FV-1200 Confocal laser-scanning microscope. Argon laser lines of 458 nm and 514 nm were used to excite mTFP1 and mVenus fluorophores, which represent the donor and the acceptor, respectively. For proper image recording, a HyD SMD detector was employed by gating a spectral acquisition window of 486-502 nm for the donor and 524-600 nm for the acceptor. FRET analysis was performed using SymPhoTime 64 software (PicoQuant) and FRET efficiency was calculated: 1-τdonor, Rac1 or Rhoa-2G /τdonor, Rac1 or Rhoa-2G-control.

### Molecular dynamics (MD) simulations

In this work, CHARMM-GUI webserver^53^, Martini coarse-grained force field^52^ and GROMACS software^76^ (version 2019.6) were used to perform all MD simulations to capture the phase separation processes of three-component lipid vesicles with different radii (10 nm and 20 nm). 1,2-dipalmitoyl-sn-glycero-3-phosphocholine (DPPC),1,2-dilinoleoyl-sn-glycero-3-phosphocholine (DUPC), and cholesterol (CHOL) with the molar ratio of 5:3:2 was adapted to construct the lipid vesicles for studying the kinetics of model membrane domains^77,78^. For all simulations, a standard 1.2 nm cutoff was applied for van der Waals interactions, and the LJ potential was shifted to zero smoothly from 0.9 to 1.2 nm to reduce the cutoff noise. For columbic potential, a 1.2 nm cutoff was used for short-range electrostatic interactions, with a smooth shift to zero from 0 to 1.2 nm. The neighbor list for nonbonded interactions was updated every 10 steps with a cut-off of 1.2 nm. Periodic boundary conditions were applied in all three dimensions. All simulations were run for 5 μs with the time step of 20 fs under the isothermal-isobaric (NPT) ensemble. Snapshots and movies were rendered by VMD^79^.

### Monte carlo (MC) simulations

We employ the Monte Carlo (MC) method to investigate the response of a discretized membrane, containing membrane domains and interacting with cortical actin via transmembrane proteins, to the pressure and membrane tension. The system Hamiltonian consists of membrane elastic energy, protein-cytoskeleton binding energy, as well as raft-raft contact energy. The system configuration evolves through three types of trial moves including vertical displacements of membrane patches, lateral translations of proteins, and lateral shifts of raft patches. These trial moves are accepted or not according to the Metropolis algorithm. We perform simulations with membrane size up to 500 nm × 500 nm, and all parameters used in the simulations are sourced from exiting literatures. For a more detailed description, see our Supplementary Materials.

### Quantification and statistical analysis

Statistical analyses for all experiments were performed using Prism (GraphPad) v5.02. Statistical data are presented as median or mean±SEM or SD. Statistical tests used and p values are specified in the figure legends. Samples in most cases were defined as the number of cells counted/examined across multiple different fields of view on the same dish/slide, and represent data from a single sample within a single experiment, which are representative of at least three additional independently conducted experiments.

### Reporting summary

Further information on research design is available in the Nature Research Reporting Summary linked to this paper.

## Data availability

All RNA-sequencing data from this study have been deposited in the Gene Expression Omnibus (https://www.ncbi.nlm.nih.gov/geo/) under accession code GSE281770. All other data in the manuscript, supplementary materials, source data and custom code are available from the corresponding author upon reasonable request. Source data are provided with this paper.

## ACKNOWLEDGMENTS

We thank Professor Dake Zhang (Beihang University) for technical assistance with RNA-seq analysis. This work was supported by the National Natural Science Foundation of China (NSFC) (12222201, 82273500, 82372750), the Beijing Natural Science Foundation (L248062), the National Key R&D Program of China (2023YFC2507000, 2017YFA0506500), Fundamental Research Funds for the Central Universities (ZG140S1971, KG16301601) and the CAMS Innovation Fund for Medical Sciences (2023-I2M-3-003 and 2022-I2M-1-008).

## AUTHOR CONTRIBUTIONS

J. D., Y.B.F., and Y.Z. designed and supervised the experiments. X.B.Z., M.T., Z.K.L., Z.G., and T.L.C. performed the experiments. M.T. and B.Q.S. performed the mouse xenograft assays. X.H.W. performed Laplace pressure assay. S.H. performed Cy5-conjugated gelatin hydrogels preparation. L.L., and X.B.L. developed the computational model. J.H.F. and S.J.W. performed patient tumor samples preparation. All the authors took part in the data analysis. X.B.Z. and J.D. interpreted the data and wrote the paper.

## SUPPLEMENTAL FIGURES

**Fig. S1.**
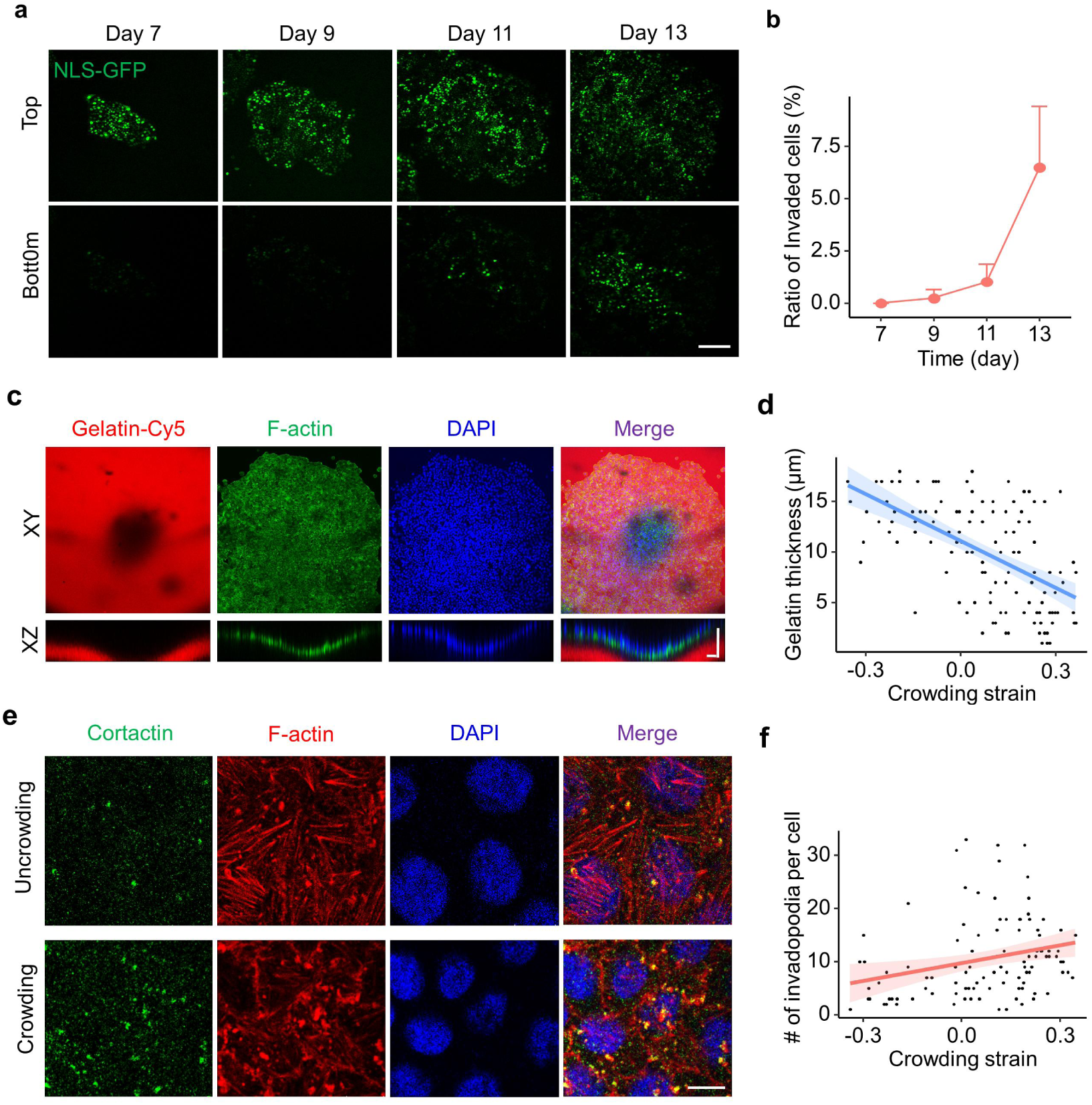
Prolonged crowding drives cancer cell invasiveness. (**a**) Transwell matrigel invasion assay in a growing monoclonal NLS-GFP^+^ HeLa cell sheets captured for 7, 9, 11, 13 days after seeding. Representative DAPI images of cells that accumulated on the top and bottom surface of the insert membranes. Scale bar: 200 μm. (**b**) The ratio of invaded NLS-GFP^+^ HeLa cells was analyzed in panel (a). (**c**) HeLa cells were plated on Cy5-conjugated gelatin hydrogels for a growing monoclonal cell sheet. F-actin was stained with phalloidin. Scale bar: 50 μm. (**d**) Quantification of correlation between Cy5-conjugated gelatin thickness underneath cells and its crowding strain in panel (c). (**e**) Representative XY slice images of cortactin and F-actin staining in uncrowded and crowded cells. Scale bar: 10 μm. (**f**) Quantification of correlation between invadopodia density per cell and corresponding crowding strain in panel (e).

**Fig. S2.**
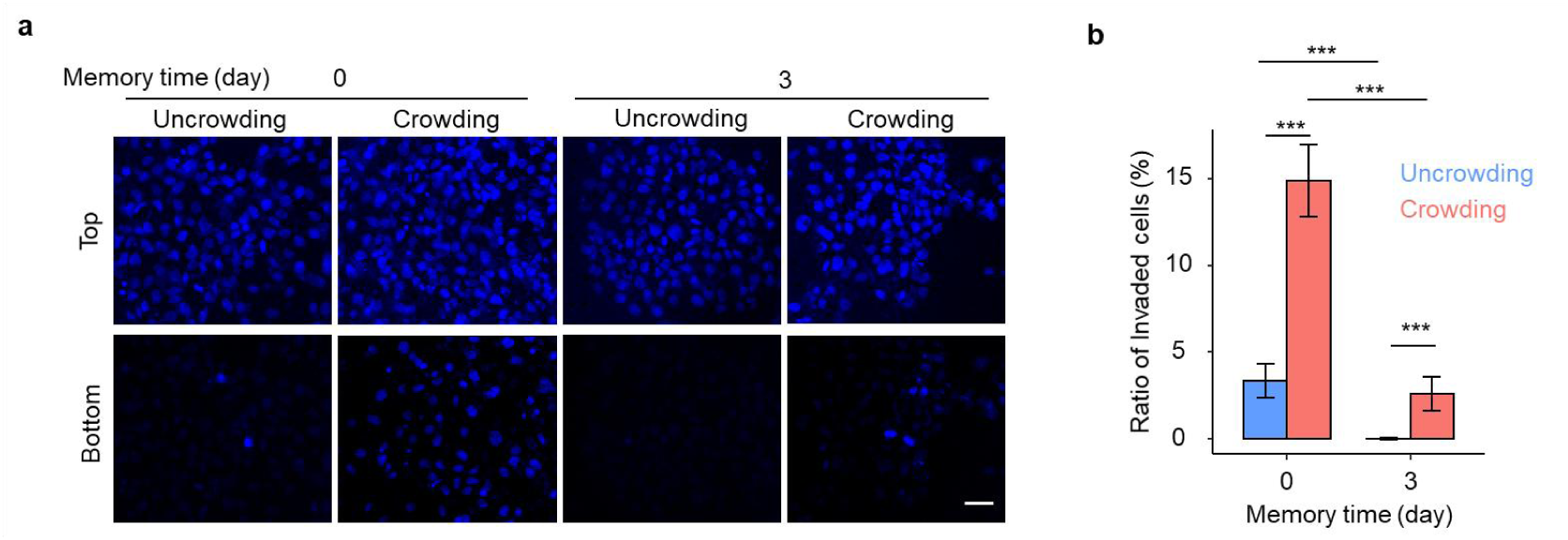
Prolonged crowding drives A431 cell invasiveness with memory retention. (**a**) Transwell matrigel invasion assay of A431 cells after uncrowding/crowding culture for 5 days and sparse culture for 0 and 3 days. Representative DAPI images of cells that accumulated on the top and bottom surface of the insert membranes. Scale bar: 50 μm. (**b**) The ratio of invaded cells in uncrowded and crowded A431 cells in panel (a). Data are presented as mean±SEM; ***p < 0.005; two-tailed unpaired t-test.

**Fig. S3.**
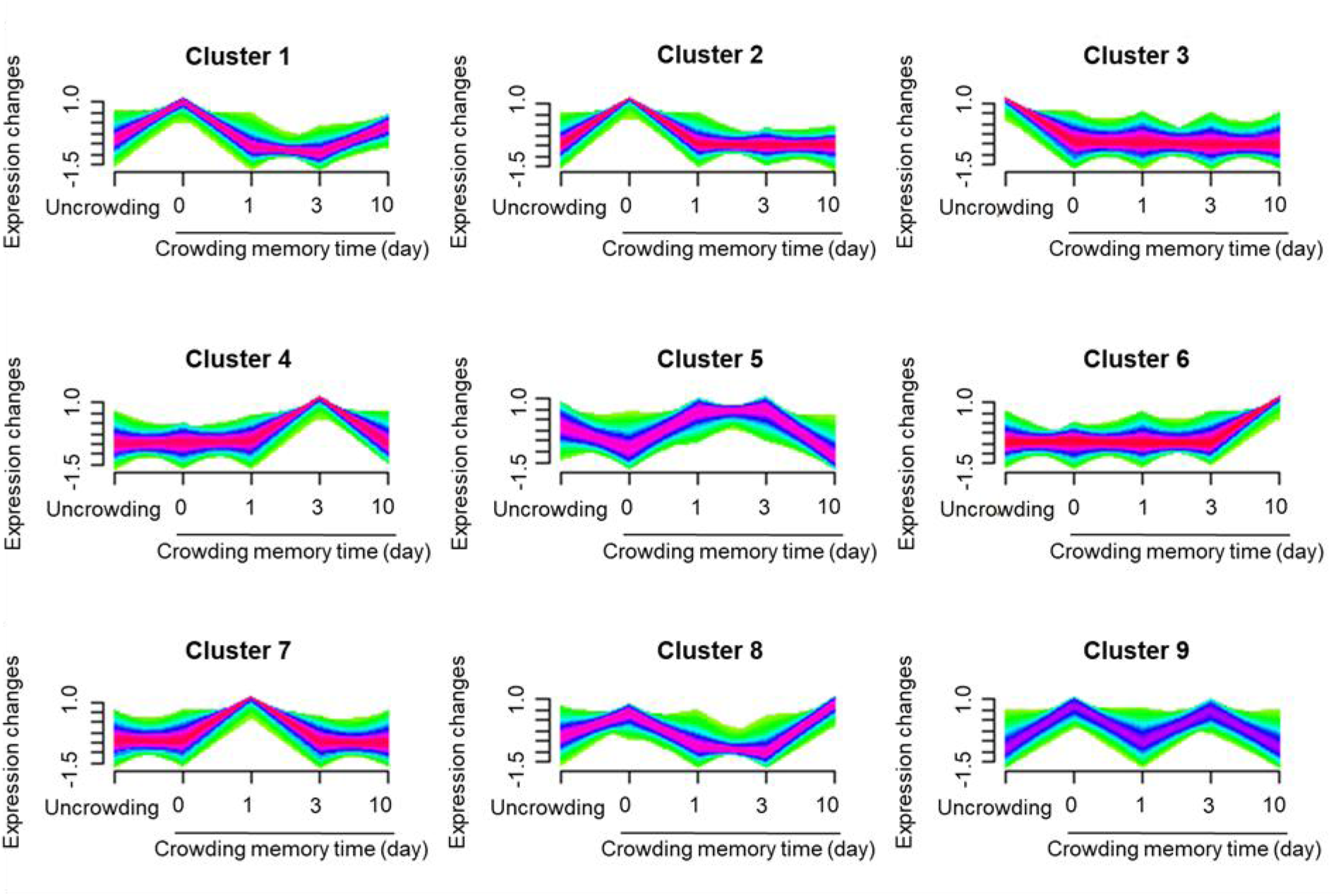
Identification of gene expression clusters in cells cultured under crowded or uncrowded conditions. Cells were primed for 20 days in crowding or uncrowding culture, followed by exposure to sparse culture for different times (0, 1, 3, or 10 days). 34,520 genes were clustered using mFuZz into significant discrete clusters.

**Fig. S4.**
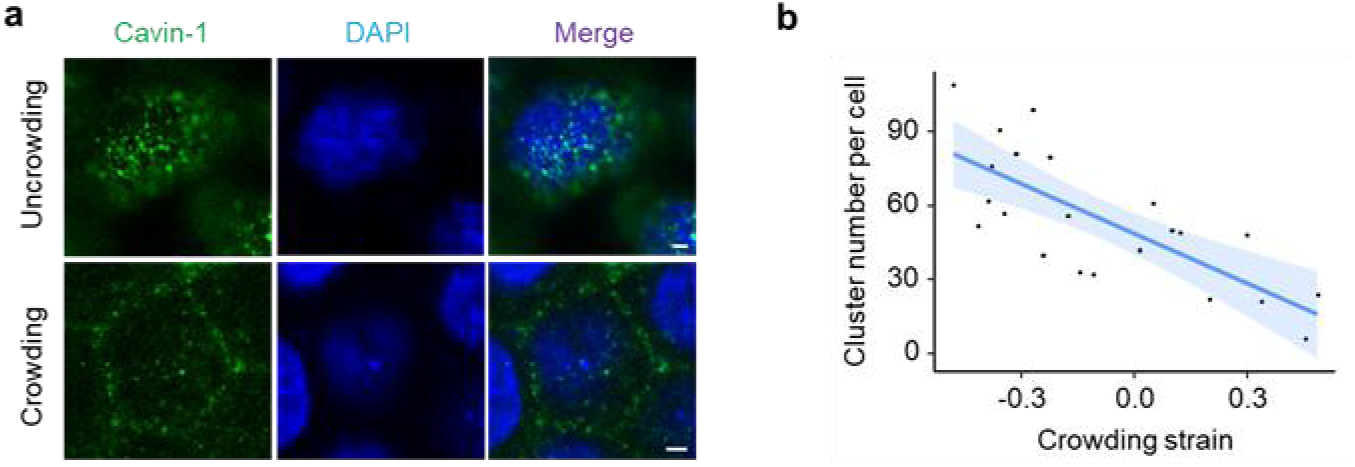
The aggregation of Cavin-1 clusters at the apical membrane of crowded and uncrowded cells. (a) Representative XY confocal images showing immunostaining for Cavin-1 at the apical membrane of uncrowded and crowded HeLa cells. Scale bar: 2 μm. (b) Quantitation of correlation between the number of Cavin-1 clusters per cell and its crowding strain.

**Fig. S5.**
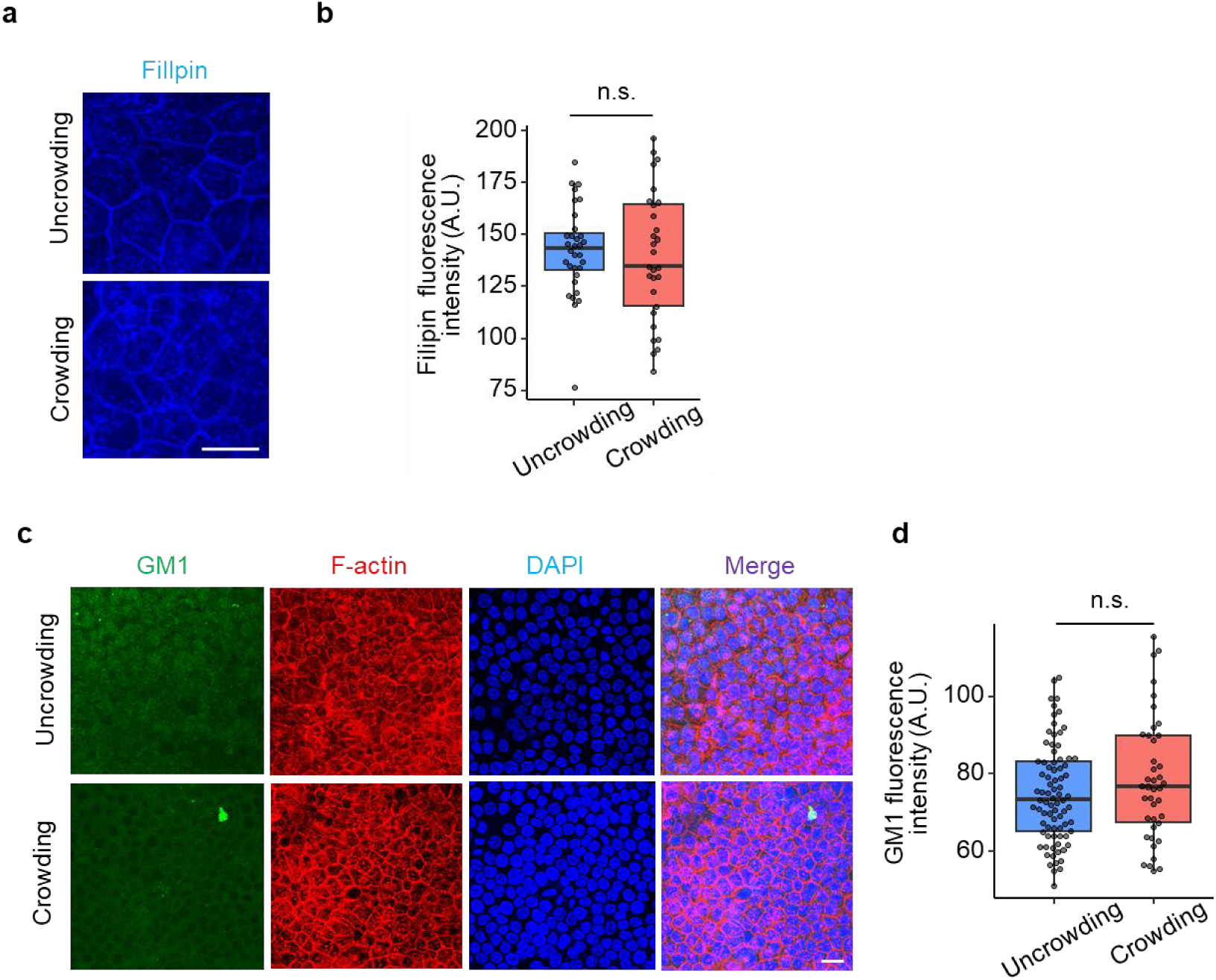
Prolonged crowding does not affect the levels of main components within membrane doamins in cancer cells. (**a**) Representative XY confocal images showing filipin staining of cholesterol in different crowding regions of monoclonal HeLa cell sheet. Scale bar: 20 μm. (**b**) Quantitation of filipin fluorescence intensity in low and high crowding HeLa cells. (**c**) Representative XY confocal images showing immunostaining for GM1 in different crowding regions of monoclonal HeLa cell sheet. Scale bar: 20 μm. (**d**) Quantitation of GM1 fluorescence intensity in uncrowded and crowded HeLa cells. Data are presented as mean ± SEM. Data are presented as n.s., not significant; two-tailed unpaired t test.

**Fig. S6.**
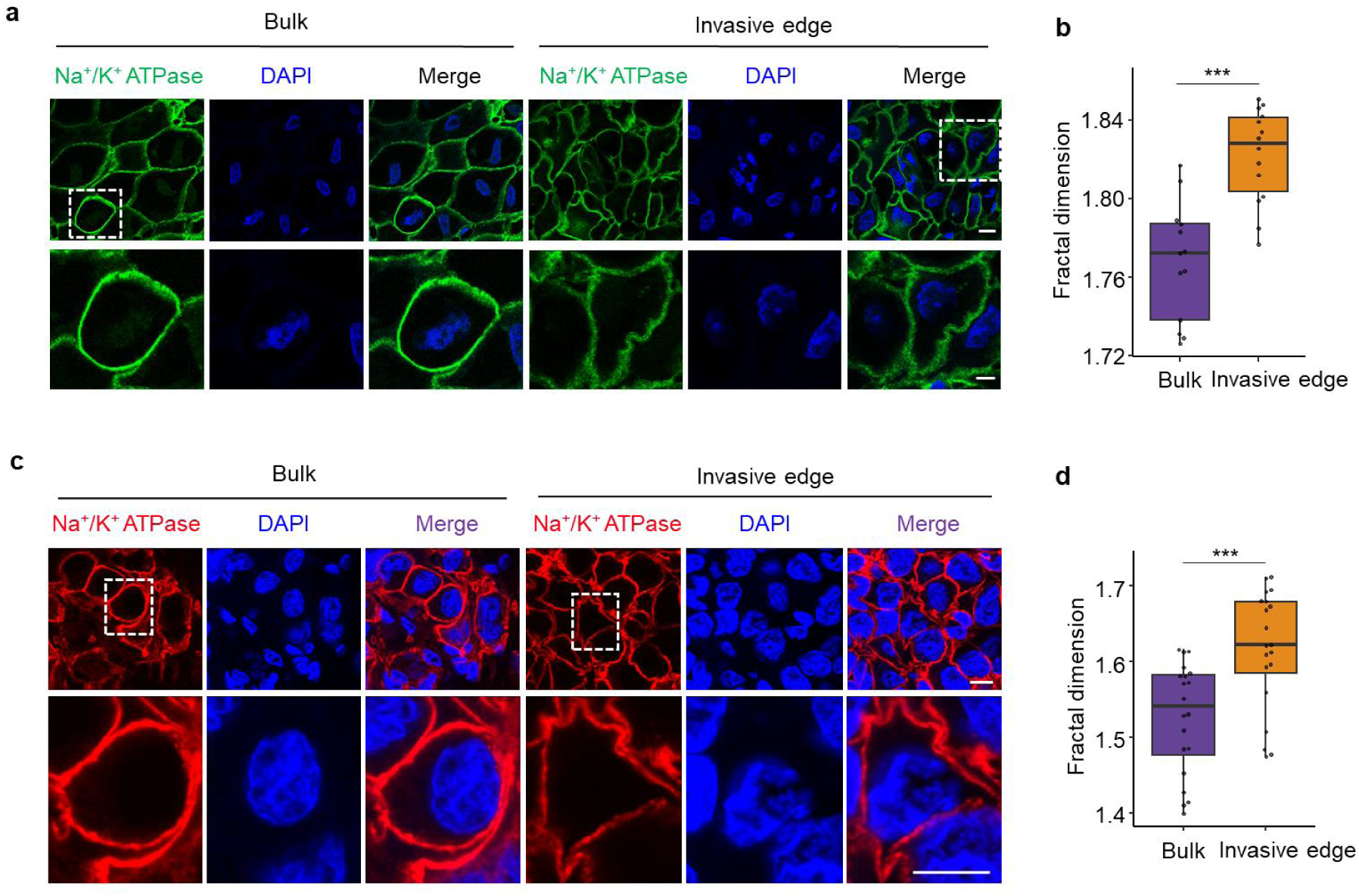
Prolonged crowding triggers a nSCTT of plasma membranes in human tumor tissues. (**a**) Representative XY slice image of sodium potassium ATPase in the bulk and invasive edge of cervical cancer tissues. Scale bar: 10 μm. (**b**) Quantitation of correlation between fractal dimension per cell and its crowding strain in panel (a). (**c**) Representative XY slice image of sodium potassium ATPase in the bulk and invasive edge of breast cancer tissues. Scale bar: 10 μm. (**d**) Quantitation of correlation between fractal dimension per cell and its crowding strain in panel (c). Data are presented as mean±SEM; ***p < 0.005; two-tailed unpaired t-test.

**Fig. S7.**
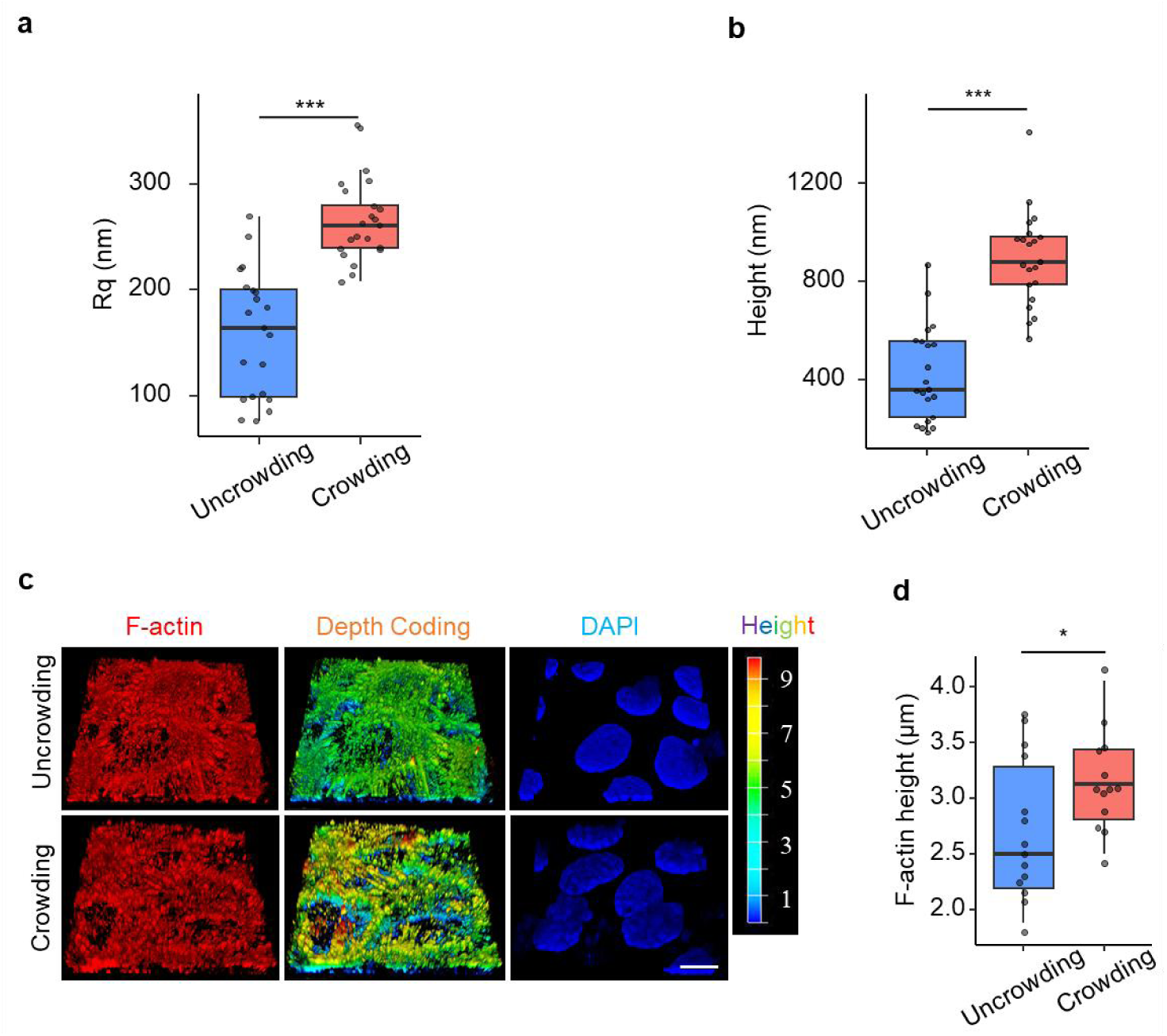
Prolonged crowding triggers membrane protrusions and cortical actin remodeling. (**a**) Root-mean-squared roughness (Rq), and (**b**) height distribution were analyzed from 3×3 μm frame ultrastructure images of HeLa cells by AFM. (**c**) Confocal 3D reconstruction of F-actin at the apical membrane of uncrowded and crowded A431 cells. Scale bar: 10 µm. (**d**) Quantification of the average height of F-actin in uncrowded and crowded A431 cells in panel (c). Data are presented as mean±SEM; *p < 0.05, ***p < 0.005; two-tailed unpaired t-test.

**Fig. S8.**
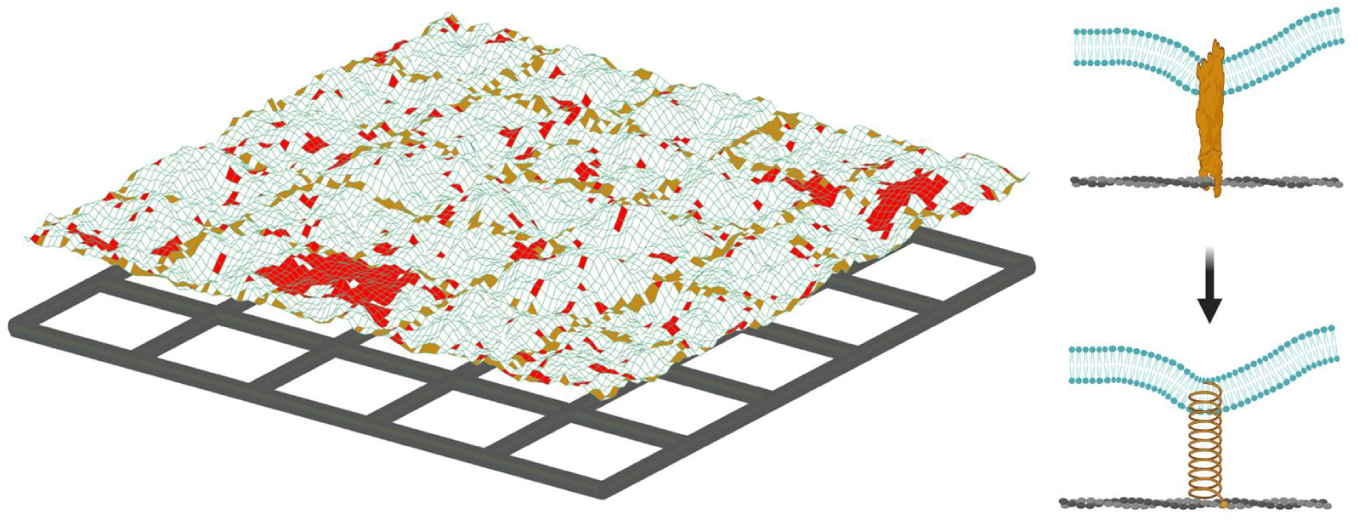
Snapshot from Monte Carlo (MC) simulations of membrane-cortical actin system. Membranes are shown in blue, cortical actin in gray, membrane domains in red. Linker proteins are indicated by square patches in yellow. The membrane-cortical actin bonds were modeled as Hookean springs.

**Fig. S9.**
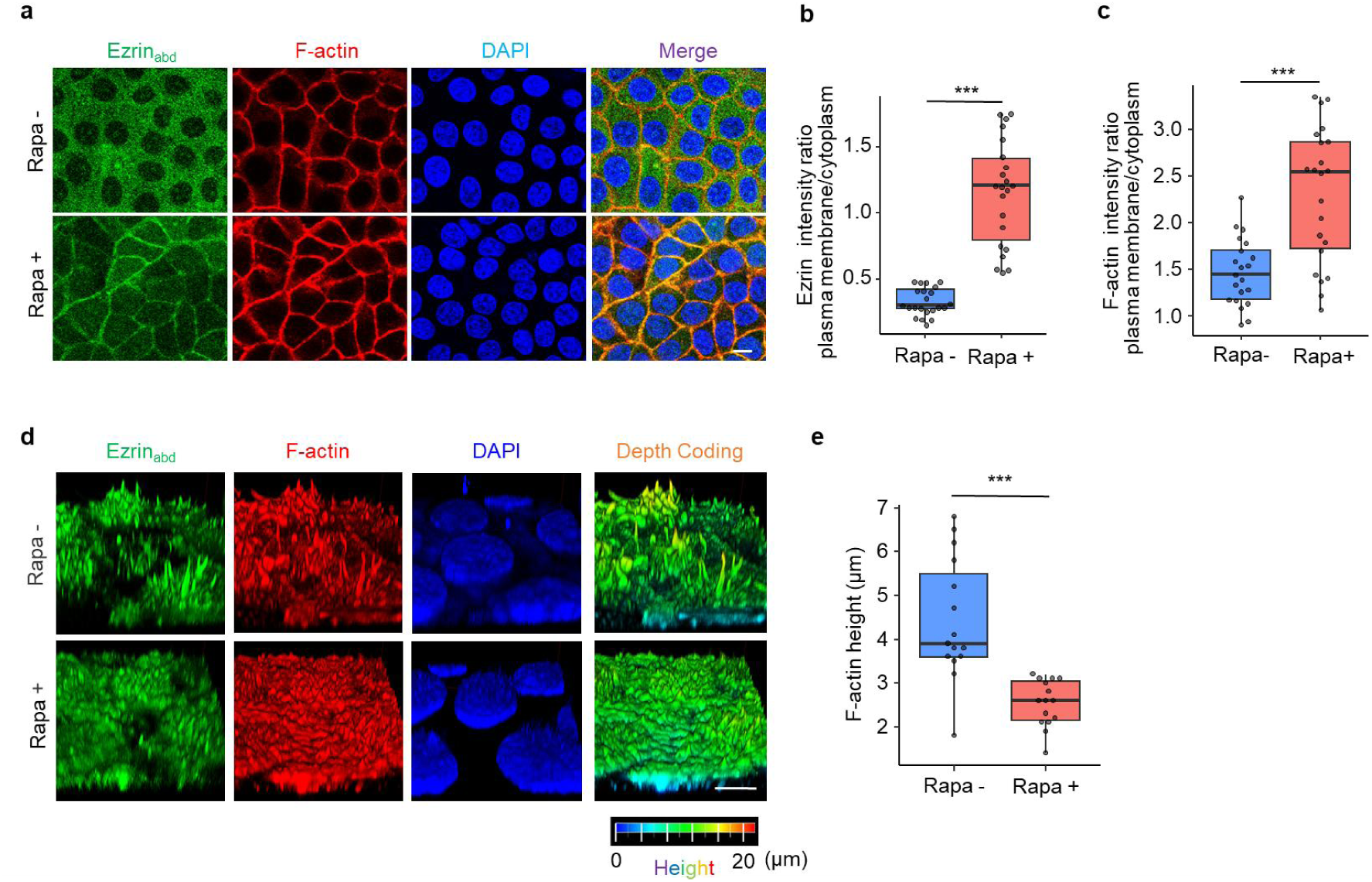
Cortical actin remodeling in crowded cells is suppressed by enhancing MCA. (**a**) Immunofluorescence analysis of Ezrin_abd_T567D in FKBP-Ezrin_abd_T567D^+^ HeLa cells treated with rapamycin or vehicle. Scale bar: 10 µm. (**b**) Quantification of Ezrin_abd_T567D intensity plasma membranes/cytoplasm ratio in panel (a). (**c**) Quantification of F-actin intensity plasma membranes/cytoplasm ratio in panel (a). (**d**) Confocal 3D reconstruction of F-actin at the apical membrane of FKBP-Ezrin_abd_T567D^+^ HeLa cells treated with rapamycin or vehicle. Scale bar: 10 µm. (**e**) Quantification of the average height of F-actin in panel (d). Data are presented as mean± SEM; ***p < 0.005; two-tailed unpaired t-test.

**Fig. S10.**
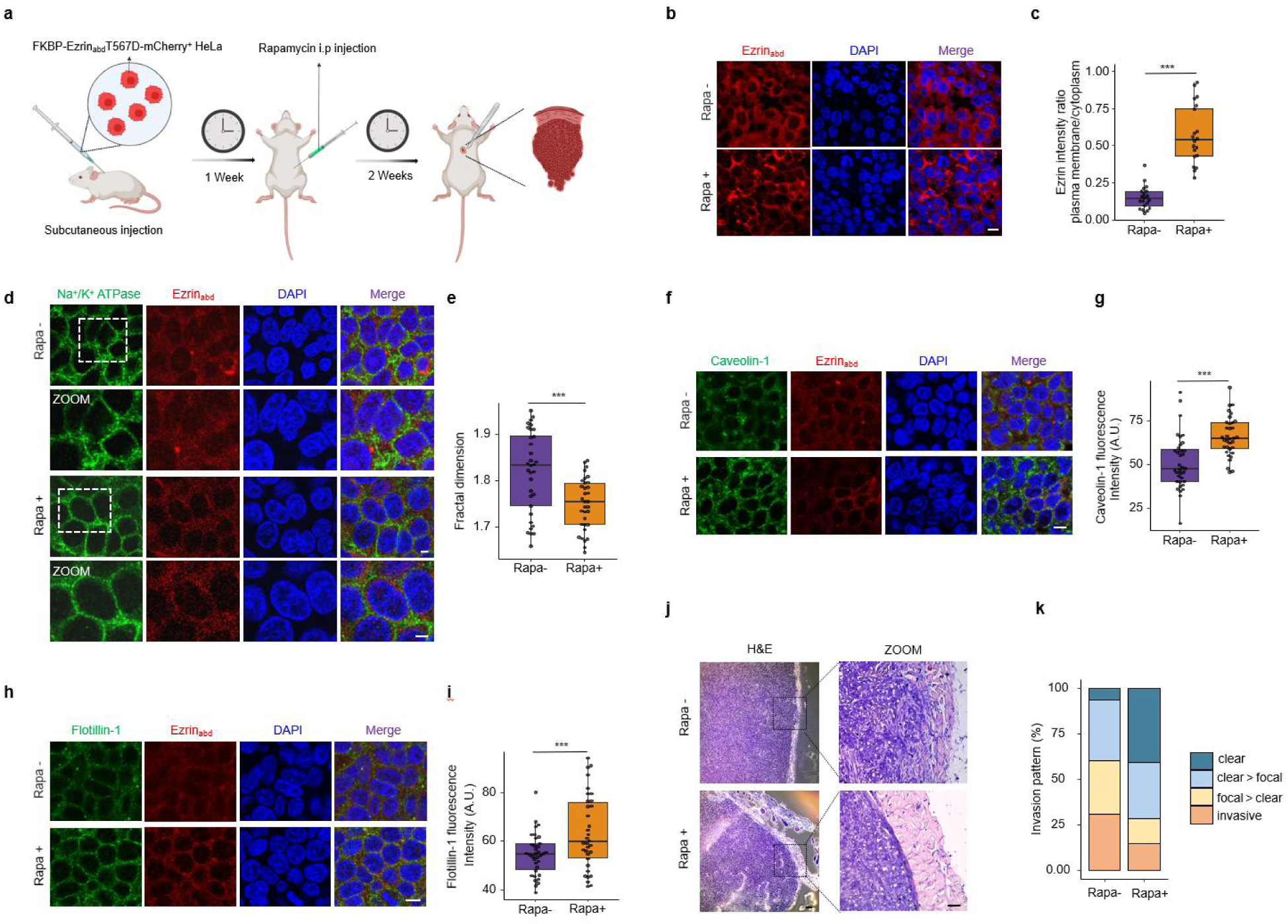
Inhibiting the nSCTT of plasma membranes suppresses cancer cell invasiveness. (**a**) Schematic diagram of subcutaneous nude mouse xenograft model using FKBP-Ezrin_abd_T567D^+^ HeLa cells. After 1 week of tumor growth, mice were treated with rapamycin or vehicle via i.p. three times per week. Tissues were harvested and tumor architectures were analyzed for the indicated parameters. (**b**) Immunofluorescence analysis of Ezrin_abd_T567D in mouse xenografts treated with rapamycin or vehicle. Scale bar: 10 μm. (**c**) Quantification of Ezrin_abd_T567D distribution in tumor cells of mouse xenografts. (**d**) Representative XY slice image of sodium potassium ATPase and Ezrin_abd_T567D in mouse xenografts treated with rapamycin or vehicle. Scale bar: 5 μm. (**e**) Quantitation of fractal dimension per cell in panel (d). (**f**) Immunofluorescence analysis of Caveolin-1 in mouse xenografts treated with rapamycin or vehicle above. Scale bar: 10 μm. (**g**) Quantification of Caveolin-1 fluorescence intensity in panel (f). (**h**) Immunofluorescence analysis of Flotillin-1 in mouse xenografts treated with rapamycin or vehicle above. Scale bar: 10 μm. (**i**) Quantification of Flotillin-1 fluorescence intensity in panel (h). (**j**) Representative images of HE staining in mouse xenografts treated with rapamycin or vehicle. Overview images, scale bar: 200 µm; Detailed images, scale bar: 50 µm. (**k**) Invasion pattern of mouse subcutaneous xenografts treated with rapamycin or vehicle. The invasion was classified as “clear”, “clear>focal”, “focal>clear”, or “invasive”. Percentages of each category are given. Only samples with sufficient surrounding muscle tissue were evaluated. Schematic image (a) was created with BioRender.com. Data are presented as mean ± SEM; ***p < 0.005; two-tailed unpaired t-test.

